# High connectivity of an iconic seabird species may buffer genetic consequences of population declines

**DOI:** 10.64898/2026.11.26.769595

**Authors:** Lucy E. Ockert, Joanna Day, John P. Y. Arnould, Christopher B. Burridge, Erin K. Clitheroe, Diane Colombelli-Négrel, Maddison J. Ledwidge, Benjamin J. Pitcher, Duncan R. Sutherland, Melanie R. Wells, William B. Sherwin, Lee A. Rollins

## Abstract

In an era of biodiversity decline, molecular techniques are increasingly used to inform conservation strategies, including delineating management units, evaluating erosion of genetic diversity, and tracing the origin of individuals. These approaches, however, rely on the ability to distinguish between populations, yet many seabird species show weak genetic structure across large ranges. Here, we generated range-wide SNP data from 772 Australian little penguins (*Eudyptula minor novaehollandiae*) at 42 colonies, sampled between 2003 and 2025, to refine estimates of connectivity, determine whether population declines have impacted genetic diversity, and to trace the origin of beach-washed individuals recovered in proximity to critically small populations. Consistent with previous microsatellite studies, overall differentiation was low (maximum *F_ST_* = 0.025) and increased with geographic distance. Spatial models indicated high gene flow across the range, except for in South Australia, where eastern and western groups delineate. Over two decades, heterozygosity and allelic richness showed no change within genetic groups, while private alleles were highly variable but appeared to be declining. All 22 beach-washed and vagrant penguins could be assigned provenance to a genetic group with 95% accuracy as determined by cross-validation testing, while assignment accuracy to colony level was 47% due to low differentiation between colonies. However, assignments indicated provenance to be distinct from stranding locations, suggesting potential dispersal rather than mortalities from nearby colonies. This work can inform conservation interventions by identifying population delineations and tracking the provenance of mortality events but also illustrates that population declines can take many generations to impact genetic diversity, particularly in highly connected species. Since genetic diversity metrics are not reliable indicators of population status, continued conservation interventions and further research into the cause of declines should be prioritised to support the long-term persistence of little penguins in Australia.

## 1. Introduction

Globally, human-induced disturbances have driven biodiversity declines across ecosystems (Ceballos et al., 2015; Storch et al., 2022). Genetic data are increasingly used to inform decisions regarding conservation interventions, including the identification of management units, to assess adaptive potential by genetic diversity, and to identify the provenance of individuals involved in dispersal, mortality events, or wildlife trafficking. Each of these applications are straightforward when populations are clearly delineated. In highly connected species, differentiation can be too weak to identify clear management units and provenance assignment can be imprecise. Many seabird species exhibit low genetic differentiation across wide distributions (Friesen et al., 2007), complicating coordinated conservation efforts. However, they are one of the most threatened vertebrate groups, with 57% of species exhibiting declines (BirdLife International, 2022). Seabirds are reliant on marine and terrestrial habitats for breeding, feeding, and migration, exposing them to a multitude of threats during critical life stages (Dias et al., 2019). Furthermore, monitoring population trends at individual colonies is often challenging due to the logistical challenges associated with accessing remote breeding habitats, meaning that extensive long-term datasets are often sparse. Effective conservation efforts rely on identifying management units and the connectivity between these units, while assessing the genetic diversity and adaptive potential within these species is central to their long-term persistence (Frankham et al., 2010; Hansen et al., 2025; Hohenlohe et al., 2021).

Declining populations become increasingly vulnerable to genetic processes such as inbreeding and genetic drift, eroding allelic variation and reducing heterozygosity, ultimately limiting adaptive capacity and long-term population persistence (Frankham et al., 2010). Where populations are genetically differentiated, the loss or reduction of individual populations can result in an irreversible loss of a species’ total genetic variation (Friesen et al., 2007; Walther et al., 2002). These risks may be heightened in range-edge populations, particularly where small population sizes are coupled with environmental conditions that are approaching a species’ physiological limits (Eckert et al., 2008; Hampe & Petit, 2005). However, genetic connectivity may buffer declining populations against genetic diversity losses by introducing new genetic variation and counteracting genetic drift (Bustillo-de la Rosa et al., 2022; Turnock et al., 2025). Detectable changes in genetic diversity also frequently lag behind population declines, particularly in species that are long-lived, have long generation times, overlapping generations, or large populations across wide distributions in which genetic drift is slowed (Charbonnel et al., 2022; Gargiulo et al., 2025). In such species, current genetic diversity may be representative of previous generations rather than recent declines (Pinto et al., 2023). This lag makes it challenging to assess the early impact of population declines using molecular markers (Hoban et al., 2014), but diversity assessments succeeding declines can provide pre-decline estimates for future comparison. Combined with estimates of genetic connectivity, such baselines can indicate whether gene flow is likely to offset drift at declining colonies, or identify any isolated populations that may continue to lose genetic diversity.

The little penguin (*Eudyptula minor*) is the smallest penguin species and the only one to breed on mainland Australia, distributed across 5,000 km of southern Australian coastline and around New Zealand (Marchant & Higgins, 1990; Reilly, 1994). Two distinct lineages have been proposed as separate species, *Eudyptula novaehollandiae* (Australia and southeast New Zealand) and *Eudyptula minor* (New Zealand) (Grosser et al., 2015; Peucker et al., 2009; Tavares & Baker, 2008). However, these taxa are currently treated as subspecies, and we therefore follow the subspecies classification here, with the Australian and southeastern New Zealand lineage referred to as *Eudyptula minor novaehollandiae*. Despite a large overall population size (estimated at ∼500,000) and being listed as ‘Least Concern’ (BirdLife International, 2020), many colonies around Australia are declining. Most extant Australian colonies now persist on offshore islands following the extirpation of many former mainland colonies (Barton, 1978; Dann, 1992; Marchant & Higgins, 1990; NSW National Parks and Wildlife Service, 2000; Stahel et al., 1987; Stevenson & Woehler, 2007), and declines have been recorded since the early 2000s in all five states across their distribution (BirdLife International, 2024; Cannell, 2025; NSW DCCEEW, 2026; Sutherland et al., 2026). Several historically large colonies have declined drastically: Granite Island (South Australia) declined from >1,500 penguins in 2001 to 26 in 2012 (Colombelli-Négrel et al., 2022); Barunguba Montague Island (New South Wales) from approximately 5000 pairs in 2000 to fewer than 300 in 2024 (NSW DCCEEW, 2026; Weerheim et al., 2003); Gabo Island (Victoria), formerly the largest colony, by ∼95% in nine years (Sutherland et al., 2026); and Penguin Island (Western Australia) by ∼95% since 2008 (Cannell, 2025). Many declines lack a clear cause, but have been linked to predators, disturbance at breeding sites, interactions with watercrafts and fisheries, and climate-mediated pressures impacting prey availability and penguin survival (Blamey et al., 2024; Cannell et al., 2016; Chiaradia et al., 2010; Colombelli-Négrel et al., 2022; Crawford et al., 2017). The recent arrival of highly pathogenic avian influenza (HPAI H5N1) in Australia (Neave et al., 2026) may further threaten vulnerable little penguin colonies. While impact of this disease on little penguins is not yet determined, extensive mortalities due to HPAI H5N1 are already evident in other seabird species (Abolnik et al., 2023; Banyard et al., 2024; Banyard et al., 2022; Clessin et al., 2025; Munoz et al., 2024). The spatial extent and magnitude of these declines highlight the need to assess whether genetic diversity has been eroded and whether cross-jurisdictional actions are required due to genetic connectivity.

Previous investigations into the population structure of little penguins in Australia have found weak genetic differentiation among colonies across most of Australia (Burridge et al., 2015; Overeem et al., 2008). The exceptions to this were colonies in eastern South Australia, where genetic structure was more pronounced, and range-edge colonies in Western Australia, which were more differentiated from the rest of the range (Burridge, 2020; Burridge et al., 2015; Colombelli-Négrel et al., 2020b; Overeem et al., 2008; Peucker et al., 2009; Vardeh et al., 2023; Wienecke, 1993). While population structure is well resolved at lower sampling density, these assessments used microsatellites and mitochondrial DNA, or single nucleotide polymorphisms (SNPs) at a finer-scale restricted to a subset of the species range. Hence, a fine-scale range-wide assessment of population structure and connectivity using next-generation sequencing is lacking. Furthermore, temporal trends in genetic diversity following population declines remain unresolved, despite the importance of understanding whether these declines have reduced the adaptive potential of little penguins as threats, such as climate variability and disease, are heightened. Additionally, beach-washed penguins were recently recovered in proximity to critically small colonies, including Granite Island (SA) and Penguin Island (WA). It is unclear whether these penguins originated from these colonies, or if they are failed attempts at dispersing, providing an opportunity to further investigate declines at threatened colonies or assess the species’ vagility.

This study provides the most detailed and wide-ranging genetic assessment of Australian little penguins to date, combining extensive fine-scale sampling across the Australian distribution of the species and high genomic resolution to assess spatial and temporal patterns of genetic variation. Specifically, we aim to: (i) determine the extent of genetic connectivity across the range to identify whether colonies are genetically distinct or part of a larger population, (ii) assess whether recent population declines have impacted genetic diversity, and (iii) determine the provenance of beach-washed penguins. These findings can be used to inform evidence-based conservation strategies aimed at maintaining genetic diversity, connectivity, and long-term resilience in little penguin populations.

## 2. Materials and Methods

### 2.1. Sample Collection and Preparation

We utilised various tissue types collected across the Australian little penguin distribution between 2003 and 2025, preserved using different approaches (Table S1). To collect contemporary blood samples (2021 – 2025) in NSW, TAS, VIC, and WA, we drew 50–200 µL of blood by venipuncture of the medial metatarsal vein using a 25-gauge needle. We collected blood samples from TAS in EDTA tubes (see Wells et al. (2024) for details) and preserved blood samples collected in NSW, VIC, and WA on Whatman® Grade 5 qualitative filter papers (Cytiva, Marlborough, Massachusetts, USA; CAT No. 1005-070) or QIAcard FTA Micro cards (Qiagen, Hilden, Germany; CAT No. WB120310), which were then left to air-dry at room temperature for up to 12 hours and stored in individual zip-lock bags at -20 °C for filter papers or room temperature for FTA cards. Details of sample collection for archived DNA can be found in Burridge et al. (2015). We also collected samples from vagrant and beach-washed little penguins, including one little penguin that landed on Ichaboe Island (Namibia, Africa) in April 2005 (see Peucker et al. (2009) for further details), five beach-washed penguins recovered in NSW between 2022 and 2024, nine beach-washed penguins recovered in SA in 2024, and seven beach-washed penguins recovered in WA in 2025. In total, this project used 822 samples of known provenance from 42 sites across Australia, and 22 samples of unknown provenance (Figure 1A, Table S1).

**Figure 1.**
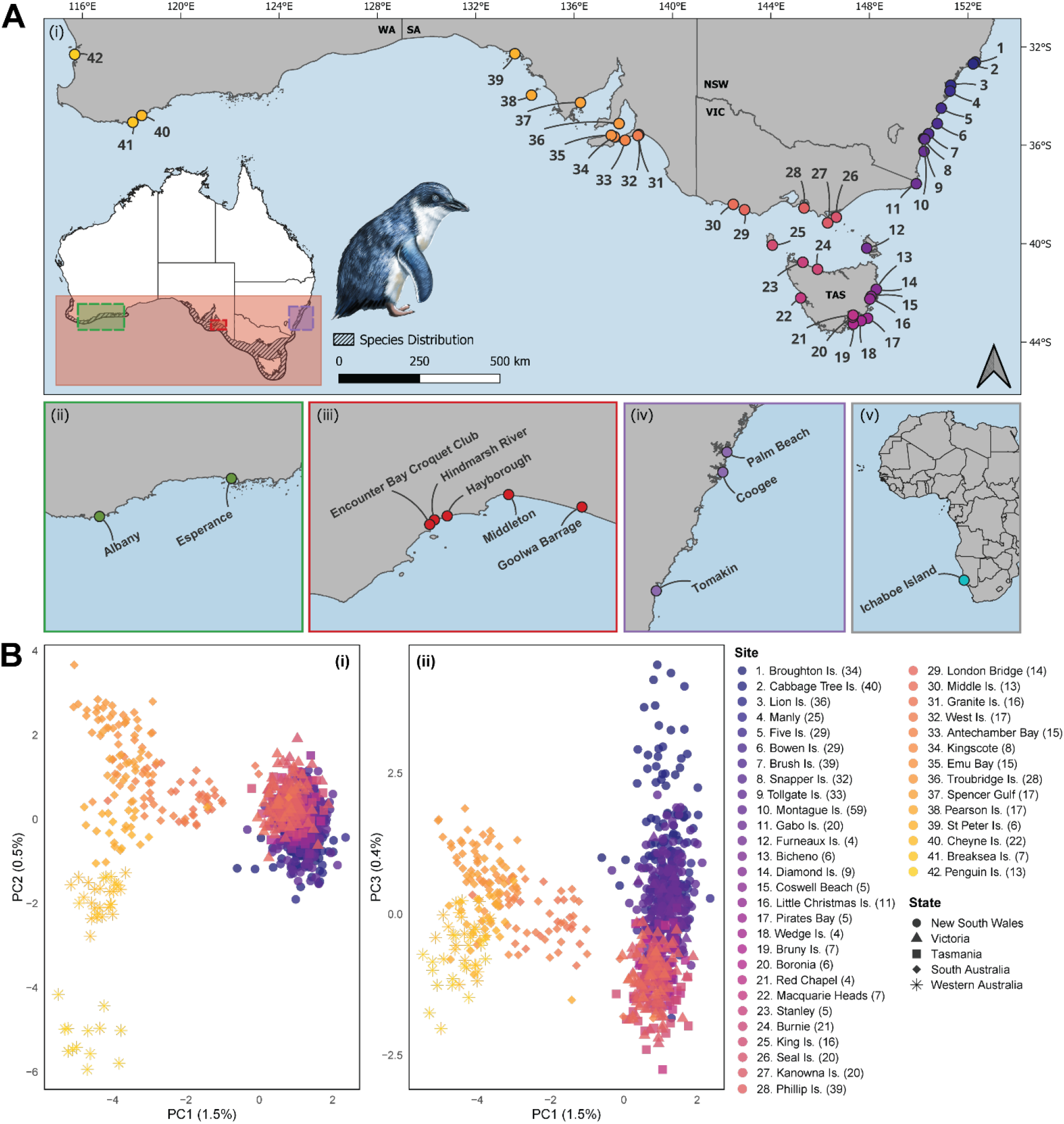
(A) Map of locations at which little penguin samples were collected, with numbers and colours corresponding to sites in (B). (i) Colonies where samples of known provenance were collected. Colour gradient represents the longitudinal position of each colony (blue = higher longitude, yellow = lower longitude). The striped polygon included in the inset map shows the species’ distribution (BirdLife International and Handbook of the Birds of the World, 2022). Additional samples were opportunistically collected from beach-washed or injured penguins of unknown provenance in (ii) Western Australia, (iii) South Australia, and (iv) New South Wales, where the outline of these maps corresponds to the coloured boxes in the inset map. A sample was also collected from a vagrant penguin on Ichaboe Island, Namibia, Africa (v). (B) Principal Coordinate Analysis (PCoA) depicting (i) PC1 and PC2, and (ii) PC1 and PC3 of 772 little penguins of known provenance, with site numbers and colours corresponding to those in (A), and final sample size (post-filtering) listed in parenthesis. Map shapefiles were sourced from Australian Bureau of Statistics (July 2021 – June 2026) Inter-Governmental Authority on Development Climate Prediction and Applications Centre (2017). Map was created using QGIS (v3.30.2-’s-Hertogenbosch) (QGIS Development Team, 2025).

To extract genomic DNA from blood in Longmire’s buffer (Table S1), we used the DNeasy Blood & Tissue Kit (Qiagen) following the manufacturers ‘Purification of Total DNA from Animal Blood or Cells’ protocol. We used approximately 15 µL of red blood cells (RBCs) suspended in Longmire’s buffer to obtain at least 1000 ng of DNA for sequencing. To increase DNA yield, we modified the protocol by (i) warming the elution buffer to 60 °C, (ii) incubating the elution buffer on the membrane for 10 min, and (iii) repeating the elution process and then combining them.

For the tissue samples (Table S1), we extracted DNA using the Puregene Kit (Qiagen), following the manufacturers ‘DNA Purification from Tissue’ protocol. For samples of blood on FTA cards or filter paper, we modified this protocol by punching approximately four 3 mm discs using a Harris UniCore Punch (Qiagen), sterilising the punch with ultraviolet light between samples to prevent cross-contamination. We increased DNA yield by (i) incubating the discs in lysis buffer and proteinase K at 55 °C with shaking at 350 rpm overnight on a thermoshaker, (ii) incubating samples with protein precipitation solution on ice for 15 min, before centrifuging samples for 5 min at 15,000 x g and repeating this step twice, (iii) adding 0.5 µL (10 µg) of glycogen solution (Qiagen) to the isopropanol, (iv) incubating the isopropanol/supernatant solution on ice for 15 min before inverting the microcentrifuge tube 50 times and centrifuging the samples for 5 min at 15,000 x g, and (v) using chilled 70% ethanol to wash the DNA pellet. DNA extraction methods for the archived DNA can be found in Burridge et al. (2015). We assessed the concentration of all DNA extractions using the Qubit 1X dsDNA Broad Range Assay Kit on a Qubit Fluorometer 3 (ThermoFisher Scientific, Waltham, Massachusetts, USA) and normalised extractions for sequencing to a maximum concentration of 60 ng/µL.

When greater than 30 samples per time period were available from individual colonies, we conducted molecular sexing via a polymerase chain reaction (PCR) to sequence roughly equal proportions of male and female samples from each site and reduce sex-based bias in downstream analysis. We amplified the CHD1 gene on the sex chromosomes using the PL/PR primer pair which was specifically designed for the molecular sexing of penguins (PL: 5′-CCC AAG GAT GAT AAA TTG TGC-3′, PR: 5′-CAC TTC CAT TAA AGC TGA TCT GG-3′) (Zhang et al., 2013) and had previously been used to sex Australian little penguins (Povah, 2021). We performed PCR in 10 µL reactions consisting of 1X of GeneAmp PCR Buffer II (Applied Biosystems, Waltham, Massachusetts, USA), 2 mM MgCl_2_ (Applied Biosystems), 2 mM dNTP Mix (ThermoFisher Scientific), 0.5 U of AmpliTaq DNA Polymerase (Applied Biosystems), 0.5 µM of each PL/PR primer (Integrated DNA Technologies, Coralville, Iowa, USA), and 5–25 ng of template DNA. We ran PCRs on a Mastercycler X50i thermal cycler (Eppendorf, Hamburg, Germany) under the following conditions: 5 min at 94 °C for initial denaturation; 32 cycles of 30 s denaturation at 94 °C, 45 s of annealing at 48 °C, and 45 s of extension at 72 °C; followed by 10 min at 72 °C for the final extension and holding at 10 °C until further processing. We visualised PCR products by electrophoresis at 70 V for 2.5 h in a 2.5% (w/v) agarose gel stained with 1X GelRed Nucleic Acid Stain (Biotium, California, USA) in 0.5X TBE buffer, alongside one male positive control, one female positive control, one negative control, and 1X of GeneRuler 100 bp DNA ladder (ThermoFisher Scientific).

### 2.2. Sequencing

We sent DNA extracts across four batches to Diversity Arrays Technology (DArT; Canberra, ACT, Australia) for medium density DArTseq, a form of reduced representation sequencing (RRS) that is robust to low concentrations of DNA from a variety of tissue types, preservation methods, and DNA extraction methods (Sansaloni et al., 2011). Duplicate samples for the same five individuals were included in each sequencing order to facilitate testing for batch effects and sequencing consistency, while DArT randomly re-sequenced 153 samples as technical replicates. DArT digested genomic DNA with *PstI* and *SphI* restriction enzymes and ligated fragments to sequencing adaptors including an Illumina flowcell attachment and barcode sequences, following methods adapted from Sansaloni et al. (2011) and Elshire et al. (2011). They selectively amplified libraries in 30 PCR cycles under the following conditions: 94°C for 1 min, followed by 30 cycles of 94°C for 20 sec, 58°C for 30 sec, and 72°C for 45 sec, with a final extension at 72°C for 7 min. They pooled and sequenced equimolar amounts of amplified product as 100-cycle single-end reads on the NovaSeq X+ platform (Illumina, San Diego, USA). We processed and analysed the raw FASTQ files for downstream analysis.

### 2.3. Sequence Processing, Alignment, and Variant Calling

To process the raw data, we first assessed the length and quality of the raw DArTseq reads using FastQC (v0.11.9) (Andrews, 2019) and MultiQC (v1.13) (Ewels et al., 2016). We then ran Stacks (v2.53) (Catchen et al., 2013; Rochette et al., 2019) *process_radtags* to remove barcodes (*- b*), remove reads with uncalled bases (*-c*), discard reads with low-quality scores (*-q*), and rescue reads with single base barcode and radtag mismatches (*-r*). We reassessed quality as above and used fastp (v0.23.4) (Chen et al., 2018) to: trim the first 15 bases, including the *PstI* (TGCAG) cut site (*-f 15*); treat bases with Phred < 30 as low-quality for read-level filtering (*-q 30*); discard reads shorter than 60 bp after trimming (*-l 60*); trim the -3’ end to a maximum of 100 bp (*-b 100*); and remove excess polyG tails (*-g*).

Two scaffold-level reference genomes are available for *E. minor* (DNA Zoo, 2024; Pan et al., 2019) and two chromosome-level reference genomes are available for species within the genus *Spheniscus* (Rhie et al., 2021), a sister clade to *Eudyptula*. While using a species-specific reference genome is often best-practice, a chromosome-level assembly facilitates more accurate alignment and robust chromosome-scale analyses. Given the high degree of genome conservation among birds (Zhang et al., 2014), particularly within Spheniscidae (Pan et al., 2019), mapping to a closely related reference genome is a reasonable alternative (Bohling, 2020; Shafer et al., 2017) and was conducted here. We used the Burrows-Wheeler Aligner tool (BWA; v0.7.17) (Li, 2013; Li & Durbin, 2009) to index all four genome assemblies. Prior to alignment, we compared mapping rates from *BWA-MEM* (designed for reads from 70 bp – 1 Mbp) and *BWA-ALN* (designed for reads up to 100 bp) against all four genomes using a subset of 50 representative individuals (Table S2). We subsequently aligned reads to the *Spheniscus humboldti* assembly (GenBank Assembly #GCA_027474245.1) (Rhie et al., 2021) using *BWA-MEM,* because this algorithm aligned a greater proportion of reads than *BWA-ALN* and is more appropriate for our read lengths (60 – 100 bp). The chromosome-level *S. humboldti* assembly was selected due to its clearer chromosomal structure, minimal reduction in mapping rate relative to the species-specific references, and inclusion of both sex chromosomes due to its female source. We ran *BWA-MEM* under default parameters.

Prior to variant calling, we compared two commonly used variant calling pipelines, BCFtools (v1.20) (Danecek et al., 2021) *mpileup*/*call* and Stacks *gstacks*/*populations*, using the same subset of 50 individuals aligned to the *S. humboldti* genome with *BWA-MEM*. We selected the Stacks pipeline to call variants across the whole dataset due to its optimisation for reduced-representation data, recommended use with non-model species that have a reference genome (Wright et al., 2019), and calling of a dataset with slightly more SNPs and higher read depth but comparable missingness to the BCFtools pipeline (Table S3).

We called initial SNPs using the Stacks *gstacks* module and then ran the *populations* module specifying to drop loci that were not present in a minimum of 30% of individuals in a population (*-r 0.3*) and retain only one random SNP per tag (*--write-random-snp*). Since some sampling sites consisted of small sample sizes, the population map was specified using broad geographic groupings and rough genetic clustering as visualised in an initial Principal Component Analysis (PCA; Figure S1), with population groups consisting of: (1) individuals from the eastern range edge in NSW (Cabbage Tree Island and Broughton Island); (2) the remaining individuals from NSW, and all of those from VIC and TAS; (3), individuals from SA; and (4) individuals from WA.

We also conducted an exploratory analysis to determine whether the number of raw reads generated, the subsequent number of filtered and trimmed reads that successfully mapped to the reference genome, the average read depth per individual, and the proportion of missingness per individual were impacted by the quantity of DNA sent for sequencing (ng), sample type, preservation method, or DNA extraction method (Figure S2).

### 2.4. Variant Filtering

To determine the most suitable filtering thresholds for our dataset and to check sequencing consistency between batches, we imported the VCF into R using vcfR (1.16.0) (Knaus & Grunwald, 2017) and ran a custom R script developed by Wright et al. (2019) to calculate DArT quality metrics. We filtered to retain sites with read depths 6–26X, before assessing sequencing consistency by calculating reproducibility and relatedness between replicate samples (Figure S3). Since no large batch effects were observed, we then merged replicate samples (plate replicates and technical replicates) to reduce genotype missingness. To filter the merged dataset, we removed sites with ≥ 70% difference in coverage difference between reference and alternate alleles, ≤ 90% reproducibility, and ≥ 70% heterozygosity. We then removed loci found on the sex-chromosomes, identified and removed duplicate individuals (genotype similarity ≥ 99%), and individuals with ≥ 50% missing genotypes. To avoid the inclusion of highly related individuals biasing analyses, we identified and removed first degree relatives using the A_JK_ statistic (Yang et al., 2010) implemented in VCFtools (v0.1.14) (Danecek et al., 2011), applying a threshold of relatedness ≥ 0.4 to account for variance in full-sibling relatedness (Speed & Balding, 2015). We compared missingness and minor allele count (rather than minor allele frequency due to uneven sample sizes) thresholds to assess the impact on broad patterns of population structure (Figure S4) and then applied additional individual and genotype missingness thresholds tailored to different analyses, following recommendations from Hemstrom et al. (2024). The main dataset was then filtered to remove loci missing in ≥ 60% of individuals and a minor allele count ≤ 3, with other combinations stated in the methods and summarised in Table S4.

### 2.5. Population Genetic Analyses

We first assessed whether the period of sample collection impacted population structure by partitioning the dataset into samples collected prior to and during 2013, and those collected after 2013 to balance the availability of samples from different locations between time points. We then conducted a Principal Coordinates Analysis (PCoA) on each dataset using *gl.pcoa()* from dartR.base (v1.2.3) (Gruber et al., 2018). After confirming that overall population structure was not impacted by sampling period (Figure S5), we examined population structure of all penguins of known provenance using multiple methods. We first ran a PCoA on the entire dataset as above and then identified genetic clusters using ADMIXTURE (v1.3.0) (Alexander et al., 2009) by testing a range of *K* values. We used ggplot2 (v4.0.2) (Wickham, 2016) to visualise the ADMIXTURE output as stacked bar plots for the *K* values with the lowest cross validation error, and plotted pie charts of the average ancestry proportions within each colony using scatterpie (v0.2.6) (Yu, 2025). To avoid over-simplifying the dataset for downstream analyses, we organised samples into the genetic groups identified by ADMIXTURE (Northern New South Wales, NNS; East Australian Coast, EAC; Bass Strait and Tasmania, BST; Southern and Western Australia, SWA) (see Results, below).

To assess the degree of genetic divergence between the four genetic groups, we estimated pairwise *F_ST_* (Nei, 1987) and Mutual Information (*I*) (Sherwin et al., 2017) since *I* weights alleles by their proportional abundance and therefore gives more weight to rare alleles than *F_ST_* does, which is dominated by common alleles. Mutual Information was calculated using *gl.report.diversity()* from dartR, while *F_ST_* was calculated using *stamppFst()* from the StAMPP package (v1.6.3) (Pembleton et al., 2013) with 95% confidence intervals calculated using 1000 bootstraps. We also calculated *F_ST_* between all sampled colonies to determine whether differentiation was driven by geographic distance. Because penguins disperse via the marine environment rather than on land, we excluded land when calculating geographic distance between colonies. To do this, we first retrieved bathymetric data for Australia from the National Ocean and Atmospheric Administration National Centers for Environmental Information (2022) using marmap (v1.0.12) (Pante & Simon-Bouhet, 2013). We then calculated the least-cost distances (*lc.dist)* between all colonies, restricting movement to marine pathways by limiting the transition matrix (*trans.mat*) to areas at or below sea level (≤ 0 m depth). Where geographic coordinates fell on land, we shifted them to the nearest marine coordinate prior to calculating distances. We assessed isolation by distance (IBD) by testing the correlation between linearised genetic distance (*F_ST_*) and marine geographic distance (km) using a Mantel test with Spearman’s rank correlation and 1000 permutations, implemented in vegan (v2.7-2) (Oksanen et al., 2025).

To estimate dispersal between colonies, referred to here as the movement of individuals away from their natal colonies, we utilised Fast Estimation of Effective Migration Surfaces (FEEMS) (v2.0) (Marcus et al., 2021), which uses a stepping stone model to identify dispersal rates that deviate from those expected under IBD. We first created a triangular spatial grid at a resolution of 7 (∼ 55 km spacing) using the *dgearthgrid()* function in dggridR (v3.1.1) (Barnes & Sahr, 2026), cropped the grid to the study area (110–155°E, 30–45°S), and removed grid cells with centroids falling within the Natural Earth 1:10m land polygon to retain only ocean cells. We then constructed a 125 km coastal buffer around the Australian mainland and Tasmania in R using the sf package (v1.1-0) (Pebesma, 2018) to define the boundary for analysis. We generated PLINK input files using *gl2plink()* from dartR.base and conducted subsequent analyses in Python (v3.12). We imported the genotype data, discrete grid, and sampling coordinates into FEEMS and imputed missing genotypes with the mean value at each SNP using *SimpleImputer* from scikit-learn (v1.8.0) (Pedregosa et al., 2011). We selected optimal regularisation parameters (λ and λ_q) for the FEEMS model via joint 5-fold cross-validation, where λ controls the smoothness of the estimated migration surface and λ_q controls node-specific residual variances (Marcus et al., 2021). We fit the final model with the validated parameters (λ = 7.74, λ_q = 0.1) and assessed model fit by comparing fitted and empirical genetic distances using a linear regression as implemented in FEEMS.

To verify the FEEMS results and determine recent dispersal rates (2–3 generations) between the four population groupings, we used BayesAss (BA3) (v3.4.4) (Wilson & Rannala, 2003). We additionally filtered the dataset to reduce missingness (Table S4) and ran four replicate runs of BA3 with different random seeds to confirm consistent results, performing 5 million MCMC iterations (*-I 5000000*), discarding the first 1 million as burn-in (*-b 1000000*), and sampling every 1000 iterations (*-n 1000*) (Figure S6). We used default mixing parameters and assessed convergence by visually inspecting the log-probability and rate parameter trace plots. We generated a chord diagram using circlize (v0.4.17) (Gu et al., 2014), and only plotted rates that were supported by a Savage-Dickey density ratio test.

To assess genetic diversity across the species’ range, we implemented a moving window approach in wingen (v2.2.0) (Bishop et al., 2023), specifying a resistance layer to exclude non-marine demes. We calculated unbiased expected heterozygosity (*uH_E_*) using a 5 x 5 cell sliding window, rarefying to five samples to address unequal sample sizes, and masked the output using the *Eudyptula minor* Australian distribution shapefile from BirdLife International and Handbook of the Birds of the World (2022). To assess genetic diversity within and across the four genetic groups (BST, EAC, NNS, SWA), we calculated observed heterozygosity (*H_O_*) and expected heterozygosity (*H_E_*) using *gl.report.hererozygosity()* from dartR.base, using 1000 bootstraps to calculate confidence intervals. We calculated mean allelic richness (*A_R_*) using *allel.rich()* in PopGenReport (v3.1.3) and private alleles (*P_A_*) using the *private_alleles()* function in poppr (v2.9.8) (Kamvar et al., 2014), both involving a single rarefaction to the smallest sample size and 1000 bootstraps to account for uneven sampling. To examine temporal changes in genetic diversity, we implemented the same methods but grouped individuals within each region into three sampling periods, reflecting periods of sampling effort: Period A (2003–2007), Period B (2012–2015), and Period C (2017–2025). We excluded all WA samples from temporal analyses since sampling at these colonies was limited an uneven between time periods, and additionally excluded groups with fewer than 10 individuals from calculations to avoid excessive down sampling, retaining two time periods per group. When calculating *P_A_*, we used this same approach to calculate across two time periods but also repeated the analysis for three time periods where sampling availability permitted (i.e., did not exclude the groups with less than 10 samples), rarefying to the smallest sample size per time period within each group.

### 2.6. Assigning Provenance

To determine the putative provenance of the 22 beach-washed penguins, we first re-ran a PCoA as above, including the penguins of unknown provenance with those of known provenance to visualise their position relative to the genetic groups. To formally assign each individual to its most likely genetic group and colony of origin, we utilised locator (v1.2.1) (Battey et al., 2020), a supervised machine learning method that predicts geographic coordinates of samples based on their genotype. We used a reference dataset consisting of 766 penguins of know origin from 42 colonies, imputed missing genotypes from allele frequencies *(--impute_missing*), and left network parameters as default. We trained the models under 10 random seeds, using 90% of samples as training and 10% as validation, and ran 10 bootstrap replicates per seed. To determine the provenance of beach-washed penguins, we retained the most frequent assignment per sample across replicates. We re-ran the analysis, treating one sample per known colony as unknown, to assess the accuracy of assignments.

## 3. Results

### 3.1. Sequencing and Variant Calling

Reduced representation sequencing generated a total of 1,538,418,776 single-end reads across 1,010 sequence files, including technical replicates and plate replicates, resulting in a mean of 1,523,187 reads per sample (range = 742,887 – 2,268,345; SD = 235,286). After trimming, filtering, and aligning the raw reads to the reference genome, a total of 1,119,117,741 single-end reads remained, with an average of 1,108,037 reads per sample (range = 362,073 – 1,575,547; SD = 182,848). These reads aligned to the *S. humboldti* genome at a mean rate of 94.84% (range = 65.57% – 98.16%; SD = 5.69%). The Stacks pipeline called 122,573 variant sites from 1,010 samples. After filtering the VCF, merging replicate samples, removing sex-linked loci, and removing individuals that were identifying as having high genotype similarity, high missingness, or high relatedness (Figure S7), we retained 9,898 SNPs and 773 individuals of known provenance (Table S4). Additional datasets for specific analyses and their associated filtering parameters are summarised in Table S4.

Despite the large range in DNA quantity and varying sample types used for this study, we observed no clear trends in sequencing quality due to DNA quantity, sample type, preservation method, or extraction method (Figure S2). Furthermore, sequencing was consistent within and between batches, with the average relatedness between technical replicates within batches being comparable to the average relatedness between plate replicates across batches (Figure S3).

### 3.2. Population Genetic Analyses

Overall patterns of population structure were not impacted by filtering for varying call rate and minor allele count thresholds (Figure S4), and sample groupings did not differ between sampling periods (Figure S5). The PCoA identified three broad groups: (1) penguins from colonies along the east coast including those in NSW, VIC, and TAS; (2) penguins from colonies in SA; and (3) penguins from colonies in WA (Figure 1B). Within these groups, SA penguins occupied a broad region of ordination space indicating high within-group variation, and Western Australian penguins separated into two subgroups corresponding to Penguin Island and southern WA (Cheyne and Breaksea Islands), although the variance explained by a PC1 was only 1.6% (Figure 1B).

The ADMIXTURE analysis showed similar cross-validation values for *K* = 2 – 4 (Figure 2). Support was highest for *K* = 2, separating colonies into an eastern group comprised of the samples from the east coast of Australia (NSW, VIC, and TAS) and a western group comprised of samples from SA and WA colonies. The *K* = 3 plot revealed subtle structure within the eastern group, distinguishing NSW colonies from those in the Bass Strait region (VIC and TAS; hereafter BST). The *K* = 4 plot further subdivided the east coast, differentiating northern NSW colonies (Broughton Island and Cabbage Tree Island) from the remainder of NSW. Across all values of *K* = 2 – 4, colonies at Antechamber Bay (Kangaroo Island), Granite Island, and West Island in eastern SA showed admixture between eastern and western groups, while WA and western SA colonies showed uniform group membership. At *K* = 4, northern NSW colonies exhibited high proportions of a unique ancestry that diminished progressively southward along the east coast, with more southern NSW colonies showing increasing proportions of an ancestry found in BST colonies. Gabo Island, which lies on the NSW/VIC border, showed admixture between the east coast and BST colonies. Within TAS, east coast colonies carried higher proportions of Australian east coast ancestry, while the remaining BST colonies were relatively uniform in their group membership (Figure 2).

**Figure 2.**
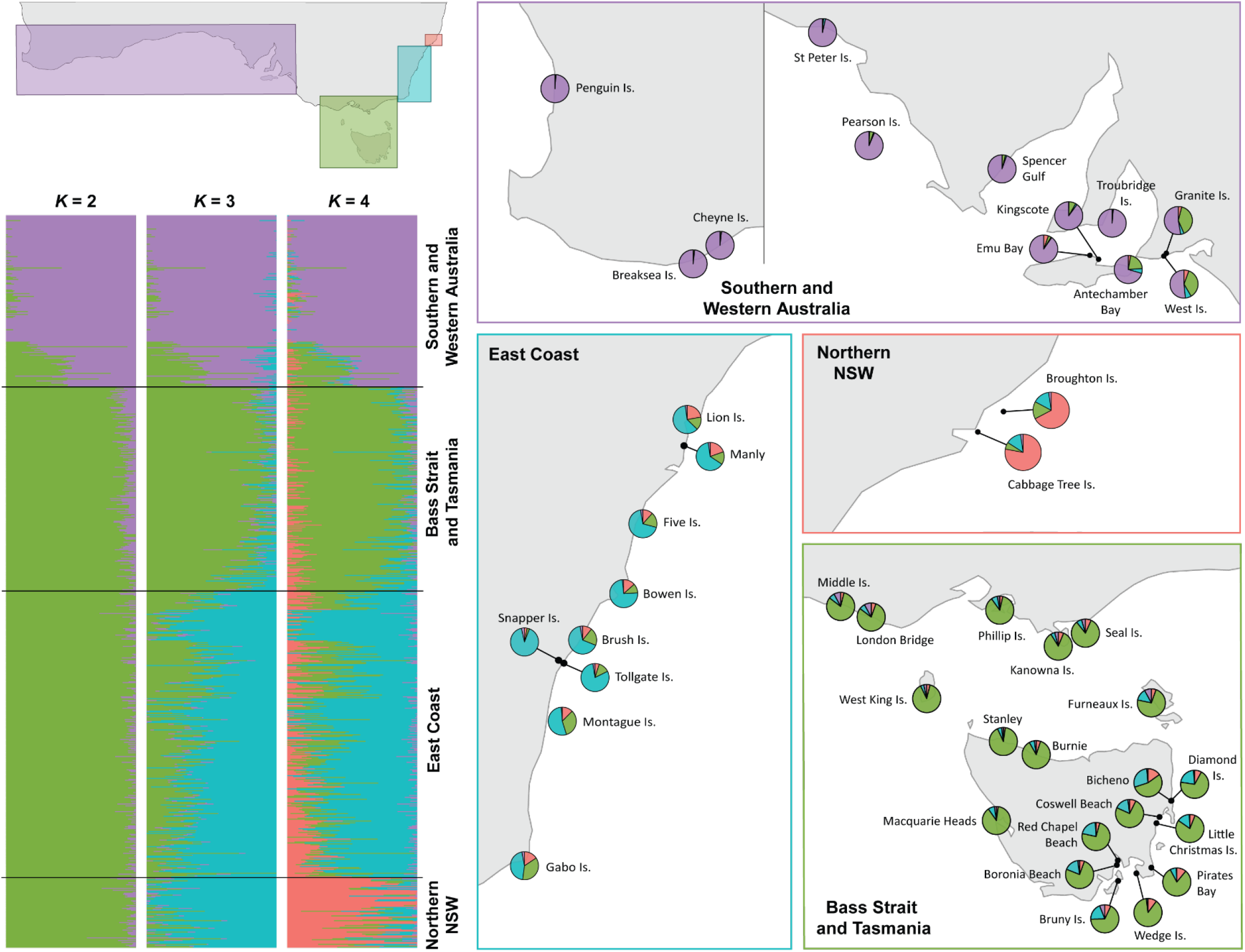
ADMIXTURE plots for *K* values ranging from 2 – 4, depicting genetic groupings of little penguins across Australia. Maps depict the average ADMIXTURE proportion at each colony, per grouping, with grouping location identified by colour (top left), with pie chart colours relating to ADMIXURE bar plot colours at *K* = 4. Individuals in the stacked bar plot are ordered based on sample location, from east to west.

Pairwise *F_ST_* values revealed low but significant differentiation between the four genetic groups (Table S5). Differentiation increased with distance, with the distal NNS and SWA groups exhibiting the highest level of differentiation (*F_ST_* = 0.0252), and EAC and BST exhibiting the lowest differentiation (*F_ST_* = 0.0032). Mutual Information showed the same pattern, ranging from 0.001 between EAC and BST to 0.010 between NNS and SWA (Table S5). However, the range of values for *I* was greater than the range for *F_ST_,* indicating that rare alleles made a small but important contribution to differentiation. The patterns of differentiation observed across the sampling range are consistent with isolation by distance, which was supported by a significant positive correlation between linearized genetic distance and marine geographic distance (Mantel *r* = 0.791, *p* < 0.001; Figure S8).

The FEEMS model fit our data well after cross-validation, with a strong correlation between fitted and empirical genetic distances (*R^2^* = 0.998; Figure S9). The estimated migration surface revealed higher levels of gene flow along the east coast of Australia and within WA, with reduced migration within SA and low connectivity between VIC and SA (Figure 3), concordant with the patterns of population structure. BA3 identified migration from EAC to all other genetic groups, with lowest gene flow to SWA, but no other migration between groups was identified (Figure 3).

**Figure 3.**
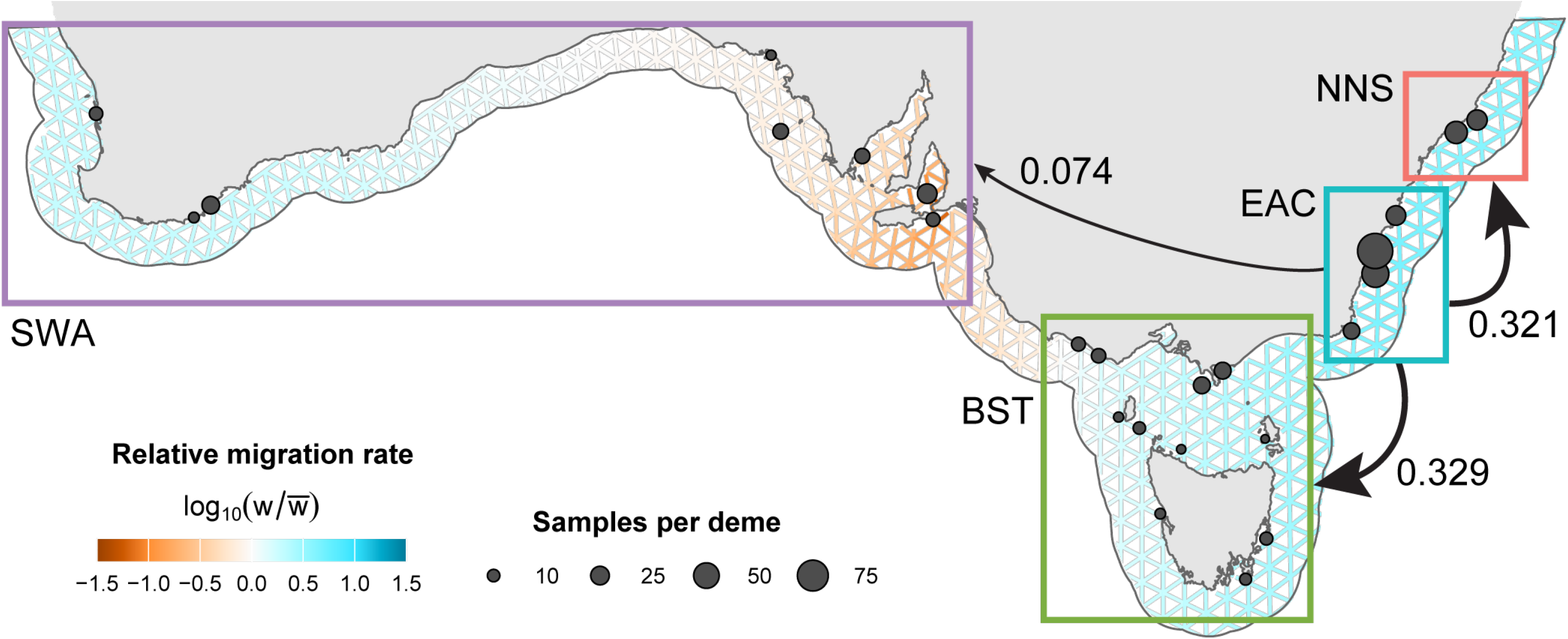
Gene flow of little penguins across Australia. Coloured boxes indicate the four genetic groups with arrows between boxes representing the migration rates (m) between these groups, calculated using BayesAss. The FEEMS output is displayed as the triangular grid around the coastline, with edge colours representing the log-scaled migration rates relative to the network wide mean where orange colours indicate gene flow that is lower than expected under isolation by distance and blue colours indicate higher gene flow that is higher than expected under isolation by distance. Node size is proportional to the number of sampled individuals within a singular deme.

Genetic diversity (*uH_E_*) was highest in colonies at the eastern edge of the sampling range and declined towards the western colonies (Figure 4A). Among the four groupings, SWA had the lowest *H_O_* (0.164) and u*H_E_* (0.1669), and highest *P_A_* (274.42), whereas NNS had the lowest *A_R_* (1.739) and *P_A_* (89.17) (Figure 4B). In each group, there were no significant changes in temporal patterns of *H_O_*, *uH_E_*, and *A_R_* across the two time periods (Figure 4C). However, at two time periods, *P_A_* declined in all groups except for NNS, where it increased (Figure 4C). When comparing three time points, a decline in *P_A_* was still evident in BST and SWA as no samples were available for the additional time point, whereas in EAC, *P_A_* was lowest at the earliest time point, and the increasing trend was still observed in NNS (Table S6).

**Figure 4.**
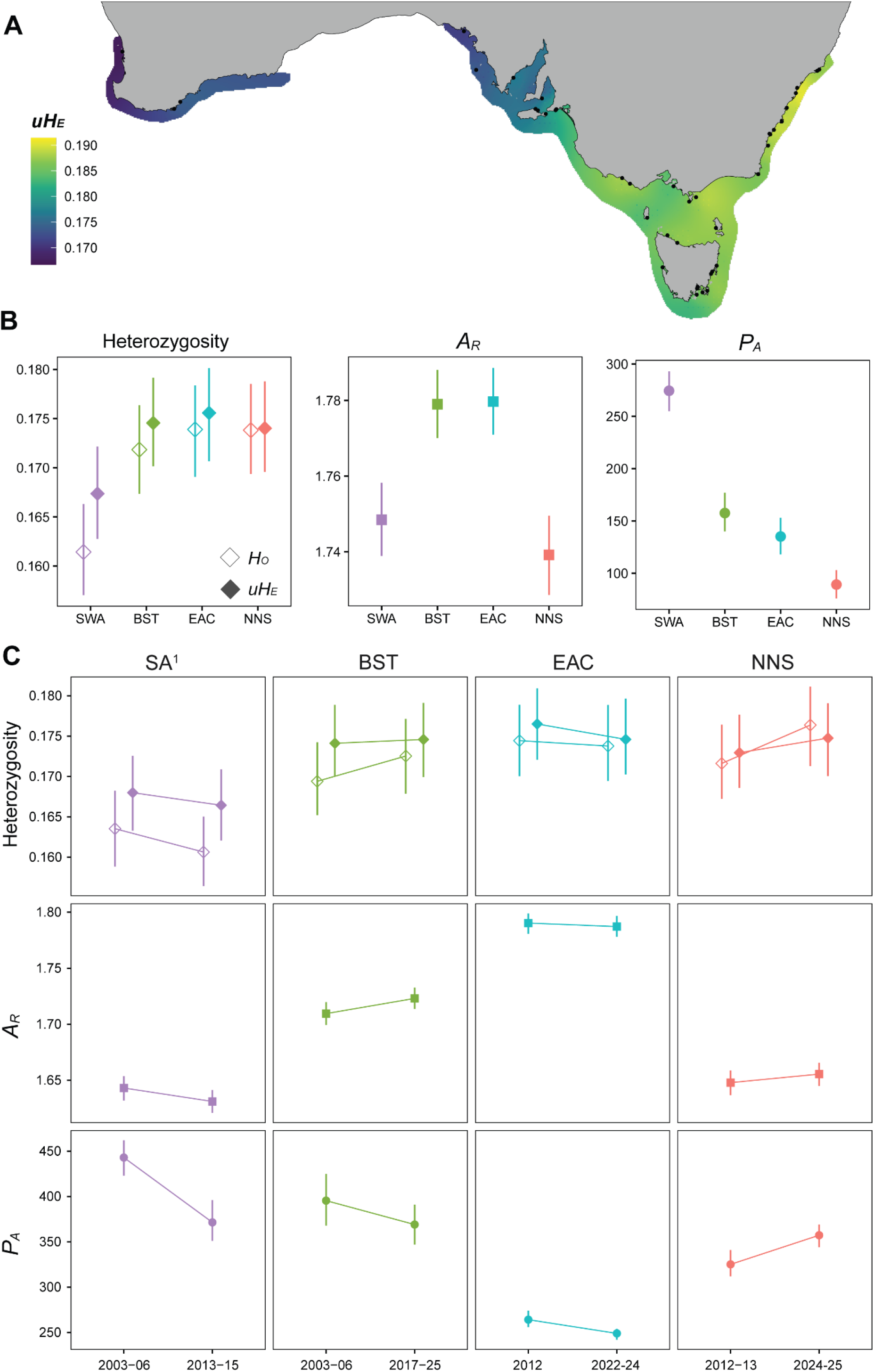
Genetic diversity of little penguins, calculated as (A) unbiased expected heterozygosity (*uH_E_*) across all sampled colonies, (B) *uH_E_*, observed heterozygosity (*H_O_*), allelic richness (*A_R_*), and private alleles (*P_A_*) per genetic grouping, and (C) same metrics as (B) but compared over two time points, where *P_A_* were calculated as those private per time period within a genetic group. ^1^Note that WA populations were excluded from temporal analyses.

### 3.3. Assigning Provenance

Based on the PCoA (Figure 5A), all beach-washed penguins recovered from WA clustered with the SWA group, while those recovered from SA, NSW, and the Namibian vagrant clustered with the eastern groups (NNS, EAC, and BST). According to locator assignments, beach-washed penguins recovered from WA likely originated from southern WA, while those recovered in SA appear to have originated mostly from BST, except for one individual that likely originated from NSW (Figure 5B). Of the five penguins recovered from NSW, two likely originated from NSW and three from BST, while the vagrant recovered from Namibia was assigned to eastern Tasmania. Cross-validation using individuals of known provenance showed that locator assigned the correct genetic group 95.4% of the time, and the correct colony 46.5% of the time, with a mean assignment error of 98.9 km (Figure S10).

**Figure 5.**
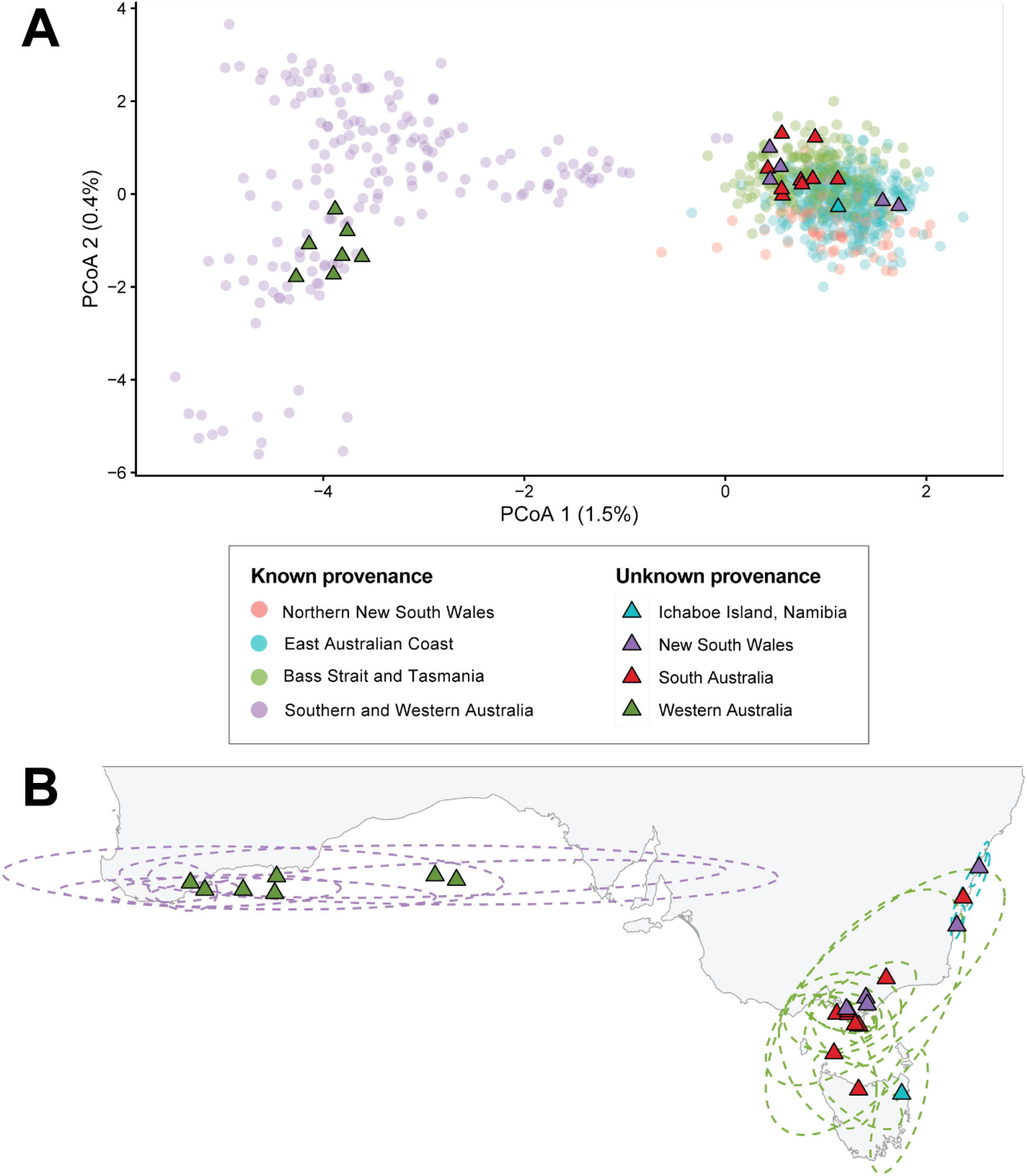
Provenance of beach-washed sampled, assessed by (A) a PCoA containing all known and unknown provenance samples, and (B) locator predictions for the origin location, based on a machine learning algorithm trained on samples of known provenance, with ellipses representing the 95% confidence intervals (based on 100 replicates; 10 bootstraps for 10 runs with different seeds) for each assignment. The legend is applicable to both plots, with all unknown provenance samples indicated in triangles coloured according to state of recovery (see Figure 1A for map).

## 4. Discussion

Here, we provide a genome-wide assessment of population structure, genetic diversity, and gene flow across the full Australian distribution of the little penguin, utilising a final dataset of 772 individuals sampled over two decades. Differentiation was consistently low and increased with geographic distance, with high gene flow within eastern and western groups that delineate near the SA-VIC border but limited gene flow between these two groups. Heterozygosity declined towards the western edge of the range, but showed no clear change over time, despite major population declines at several colonies. Beach-washed and vagrant penguins could be reliably assigned to genetic groups but less so to colonies, and provenance assignment indicated that the origin of most beach-washed birds differed to their recovery location. These findings have implications for little penguin conservation in Australia but also aid in our understanding of managing seabirds more broadly, illustrating the challenges of defining conservation units in species with continuous genetic structure, demonstrating the lag associated with detecting genetic diversity trends after population declines, and outlining the use of genetic data in tracing the provenance of stranded and vagrant birds when differentiation is low.

### 4.1. Population Structure and Connectivity

Consistent with previous studies (Burridge et al., 2015; Overeem et al., 2008; Peucker et al., 2009), we identified two main genetic groups of little penguins in Australia (eastern and western), which delineate near the SA-VIC border. Denser sampling allowed us to resolve structure at a finer resolution, particularly in NSW and TAS due to the inclusion of colonies previously not examined. We identified three subgroups in the eastern group; Northern NSW (NNS), East Australian Coast (EAC), and Bass Strait and Tasmania (BST), each containing higher proportions of their associated admixture membership that gradually declined with distance. Broughton Island and Cabbage Tree Island, the two most northern colonies in NSW, carried the highest proportions of NNS membership which progressively declined southwards, while southern NSW colonies shared increasing membership with BST. Gabo Island, on the NSW-VIC border, showed mixed membership between EAC and BST, and colonies on the east coast of TAS contained higher proportions of EAC membership than other BST colonies. This gradient along the east coast, together with high gene flow in this region estimated by FEEMS, may reflect dispersal towards the centre of the species range assisted by the southward-flowing East Australian Current (Wijeratne et al., 2018). BayesAss dispersal estimates between NNS, EAC, and BST reached the upper bound of the model (Faubet et al., 2007; Meirmans, 2014) and should therefore be interpreted as indicative of substantial gene flow rather than absolute dispersal rates. BayesAss and FEEMS dispersal estimates, partnered with low differentiation, indicate that these groups are best interpreted as segments of a continuous gradient rather than discrete populations.

The boundary between the eastern and western groups, located in eastern SA, contrasted the pattern of high gene flow observed otherwise. FEEMS estimated lower gene flow than expected under isolation by distance between VIC and SA, and the eastern SA colonies of Antechamber Bay, Granite Island, and West Island showed mixed membership of SWA and BST groups. Despite the lower gene flow in this region, dispersal between EAC and SWA was still estimated by BayesAss, and movement between VIC and SA has been documented by banding records (Dann et al., 1992; Norman et al., 2018). Although banding recoveries do not necessarily mean that these penguins are breeding and facilitating gene flow, it does indicate that the presence of a physical barrier limiting gene flow between these regions is unlikely. This pattern is instead consistent with a secondary contact between historically separated eastern and western lineages, as proposed previously (Burridge et al., 2015). While fine scale structure among SA colonies has been previously reported and debated (Burridge, 2020; Colombelli-Négrel et al., 2020a; Colombelli-Négrel et al., 2020b), our findings support reduced gene flow within SA, as expected in a contact zone, but do not identify isolated colonies.

Weak genetic structure over large distributions is common among penguins and other seabird species (Friesen et al., 2007; Munro & Burg, 2017), as demonstrated by Chinstrap (*Pygoscelis antarcticus*) (Mura-Jornet et al., 2018), Adelie (*Pygoscelis adeliae) (Clucas et al., 2014)*, emperor (*Aptenodytes forsteri*) (Younger et al., 2017), and king (*Aptenodytes patagonicus*) (Clucas et al., 2016) penguins. This low genetic differentiation may be due to recent diversification, historical admixture between lineages, and past demographic declines (Friesen et al., 2007). These processes are plausible here, given the diversification of *Eudyptula* ∼ 3 million years ago, evidence of recent introgressions between Australian and New Zealand lineages, and a steep population decline during the Last Glacial Maximum (Baker et al., 2006; Cole et al., 2022). Interestingly, the genetic structure we observed in little penguins resembles patterns reported in other southern Australian marine taxa (Bertram et al., 2023; Petrolo et al., 2021; Stiller et al., 2023) and corresponds broadly to three marine biogeographic provinces of southern Australia (Waters et al., 2010); SWA to the western Flindersian, BST to the south-eastern Maugean, and NNS and EAC correspond to the eastern Peronian province. Therefore, the low differentiation and subtle genetic structure observed among little penguins in Australia may reflect both the species evolutionary history and the historical and oceanographic processes that shape marine biogeography in this region. Given that southern Australia includes marine warming hotspots (Hobday & Pecl, 2014), understanding the connectivity and genetic diversity of marine species in this region may help predict their responses to future climatic conditions.

### 4.2. Spatial and Temporal Patterns of Genetic Diversity

We identified observed and expected heterozygosity to be highest in eastern colonies, declining towards the west, with heterozygosity lowest among SWA colonies. SWA also contained the most private alleles, consistent with its position at the western edge of the range and its lower connectivity to eastern colonies. Allelic richness was lowest at either edge of the species range in NNS and SWA, consistent with the central-marginal hypothesis which predicts reduced diversity in peripheral populations that are smaller, more isolated, or recently established (Eckert et al., 2008).

Several colonies from each genetic group have recorded steep population declines during our sampling period, however, we did not detect any changes in heterozygosity or allelic richness between sampling periods. This is perhaps unsurprising given the life history of little penguins; the generation time of ∼8 years (Bird et al., 2020) means that our sampling interval likely does not cover sufficient time for allele frequencies to shift, while records of individuals exceeding 20 years old (Dann et al., 2005) means that genetic variation from previous generations can be retained within the breeding population long after a decline occurs, particularly when generations are overlapping (Charbonnel et al., 2022; Clark et al., 2024). Furthermore, the survival of first-year birds is often quite low, with older penguins making up a larger proportion of the breeding population and exhibiting higher breeding success (Colombelli-Négrel, 2015; Sidhu et al., 2007). Together, these factors mean that contemporary diversity estimates are likely to lag demographic change (Gargiulo et al., 2025). The high gene flow observed between colonies in each genetic group may also buffer declining colonies against a loss of genetic diversity, as immigrants can reintroduce alleles lost through genetic drift (Mills & Allendorf, 1996).

In declining populations, it is predicted that rare alleles will be lost prior to measurable declines in heterozygosity (Allendorf, 1986). We detected declines in private alleles across two time periods in SWA, BST, and EAC, which may indicate an early indicator of declining genetic diversity in these regions. However, NNS demonstrated an increasing trend, and private allele trends were inconsistent when examining a third time period in EAC. Despite the use of thousands of alleles and rarefaction to account for uneven sample sizes, private alleles are sensitive to sample size and sampling error and, therefore, may not be a reliable indicator of decline. Detecting population declines early is vital for effective conservation interventions but can be hindered by genetic diversity metrics failing to reflect declines, even when extensive temporal samples are available and extreme bottlenecks have occurred (e.g. Liu et al., 2025). It is clear that population sizes at little penguin colonies have dramatically declined, although we were unable to detect a clear trend in genetic diversity here, likely due to short sampling intervals and little penguin life history traits. While continued genetic monitoring at longer intervals may be beneficial to assess genetic diversity levels, particularly if declines continue, this is not feasible for all colonies. Therefore, temporal census surveys may be more impactful at measuring population declines. Alternatively, genetic methods proposed to estimate census size directly, such as DNAdot (Sherwin, 2024), may be more reliable at detecting early declines and could greatly benefit conservation efforts.

### 4.3. Provenance of Beach-Washed and Vagrant Penguins

The fine-scale accuracy of provenance assignment of beach-washed individuals was limited due to the lack of differentiation between colonies; however, all individuals could be assigned to a genetic group. Beach-washed penguins recovered in SA and some of those recovered in NSW were assigned provenance distal to their recovery location, while others recovered in NSW and WA appear to have originated from colonies in their respective states. The seven penguins recovered from SA in proximity to Granite Island, a critically small and declining colony, appear to have originated from colonies in BST or EAC which is reassuring for the persistence of the Granite Island colony. The movement of penguins between the eastern Australian coast and South Australia has been recorded, with penguin banding studies demonstrating that juvenile little penguins from VIC will frequently disperse westwards, with recoveries as far west as Spencer Gulf, Kangaroo Island, and Long Bay (Dann et al., 1992). Additionally, non-breeding and juvenile little penguins have been shown to travel west during periods of low food availability, evidenced by multiple mortality events (Norman et al., 1992). The penguins recovered from WA showed genetic signatures consistent with SA and WA origins. Because past banding efforts were concentrated at particular colonies of interest, definitive conclusions about dispersal routes in these regions are limited; however, there are banding records of penguins travelling from VIC to WA, and from WA to SA (DCCEEW, 2025). Given these long-range movements and the westward dispersal of east coast penguins (Dann et al., 1992; Norman et al., 2018), it is plausible that SA penguins could move westwards to WA. Penguins recovered in NSW were assigned to EAC and BST, which is consistent with banding records showing that little penguins readily disperse along the east coast (Reilly & Cullen, 1982). Although limited differentiation between VIC and TAS inhibited finer-scale provenance assignment, some records also document movement of penguins between NSW and TAS (Reilly & Cullen, 1982). Interestingly, the vagrant from Namibia most likely originated from the east coast of Tasmania, despite the closer proximity of Namibia to more western colonies. Our use of SNP data enabled assignment to a higher resolution than previous mitochondrial analyses that identified this vagrant as being most closely associated with the Australian lineage (Peucker et al., 2009). Records of vagrant little penguins are sparse, though multiple sightings have been documented in Chile (Barros et al., 2015; Brito, 1999; Wilson et al., 2000), and vagrancy is well established across penguin taxa more broadly (Miskelly et al., 2012; Shepherd et al., 2021; Woehler, 2026). Overall, the low levels of differentiation and high connectivity between little penguin colonies indicate that genetic techniques can broadly identify the area of origin but cannot always accurately assign provenance to specific colonies; a limitation that has also been demonstrated for other penguin species (Roeder et al., 2001).

### 4.4. Conservation and Broader Implications

The continuous genetic structure demonstrated by little penguins in Australia supports the management of this species at a regional scale, coordinated across state boundaries. We identified four genetic groups that indicate where genetic composition varies most, however, the high gene flow between colonies indicates that these are not independent units. Colonies of high conservation concern, such as Manly in NSW, belong to a larger genetic group and does not contain genetic variation dissimilar to neighbouring colonies. Despite showing little genetic differentiation, small colonies may remain important because of their ecological significance within local ecosystems, contribution to regional population resilience, or role in maintaining the species’ distribution. Penguin Island in WA, which similarly is recognised as being of high conservation concern, represents the range-edge of the species. The western edge of the species range had elevated differentiation compared to the rest of the range, and population viability analyses have predicted higher extirpation risk of range-edge colonies in WA and in NSW (Vardeh, 2016). As such, continued demographic and genetic monitoring of this colony as well as a more extensive evaluation of the gene flow and differentiation within WA would be beneficial for the continued conservation of this colony.

Encouragingly, previous efforts of little penguin conservation demonstrate that population recovery is achievable. The Phillip Island colony, which experienced significant declines during the 1980s, is now the largest in Australia following extensive efforts to reduce land-based threats (Dann, 1992; Sutherland & Dann, 2014), while effective predator management has minimised declines at multiple additional sites (Overeem & Wallis, 2007; Rout et al., 2014; Wallis et al., 2017). Since adult survival affects colony viability more than juvenile survival (Colombelli-Négrel, 2015; Sidhu et al., 2007), conservation efforts should focus on identifying and mitigating threats that are most impactful for reducing adult mortality. However, the presence of disease (Campbell et al., 2022; Cannell et al., 2013; Harrigan, 1992), including HPAI H5N1, pose additional challenges for conservation management. Supporting breeding colonies by controlling predators and providing supplemental habitat in the form of nestboxes may improve the resilience of colonies but is unable to directly mitigate disease transmission. As such, the recent detection of HPAI H5N1 on mainland Australia (Neave et al., 2026) emphasises the importance of continued colony monitoring to facilitate the early detection of positive cases, tracking transmission pathways, and quantifying the impact on little penguins. In the event of further population declines, due to disease or otherwise, our dataset serves as a baseline to compare future genetic diversity estimates. Although we were unable to detect a definitive temporal decline in genetic diversity, the decline in private alleles among three of the genetic groups may be an early signal of genetic diversity loss due to population declines. The magnitude of recent population declines at some colonies may continue to manifest as reduced genetic diversity over longer time scales, and therefore, the continued sampling and monitoring of these colonies over multiple generations would be beneficial to assess longer-term impacts, inform conservation interventions, and support uplisting the species conservation status. Future research into the impacts of HPAI H5N1 on this species and exploring whether range-edge colonies are locally adapted to more variable climatic conditions would be beneficial for understanding the impact of threatening processes and assessing adaptive capacity in a continually changing climate.

## Acknowledgements

Thank you to Karen Gray and the Australian Museum, Nicholas Carlile, Holly Raudino, Rebecca Overeem, Amanda Peucker, Stephen Hedges, Arisha Silverlake, Lewis Vaughan, and Sandra Vardeh for the donation of samples. Thank you to Katarina Stuart for input regarding bioinformatic analysis; David Slip, Gabrielle Potts-Todd, and Lindsay Wright for their involvement in fieldwork; and Marta Stojnic, Ella Trueman, Alena Cheng and Rafaela Gurgel for assistance in the lab. This research includes computations using the computational cluster Katana supported by Research Technology Services at UNSW Sydney (https://doi.org/10.26190/669x-a286).

## Funding Statement

Funding for this project was provided by a NSW Environmental Trust grant awarded to Taronga Conservation Society Australia as part of the NSW Department of Climate Change, Energy, the Environment, and Water’s Seabirds to Seascapes project; the Holsworth Wildlife Research Endowment Equity Trustees Charitable Foundation and the Ecological Society of Australia; and the Threatened Species Initiative Consortium (https://doi.org/10.25953/3eje-q240), supported by funding from BioPlatforms Australia and enabled by the Commonwealth Government National Collaborative Research Infrastructure Strategy (NCRIS) in partnership with the University of Sydney, the Australian Government Department of Climate Change, Energy, the Environment and Water, WA Department of Biodiversity, Conservation & Attractions, and Amazon Web Services. LEO’s scholarship was provided by an Australian Government Research Training Program (RTP) Scholarship (doi.org/10.82133/C42F-K220).

## Conflict of Interest Disclosure

The authors declare no conflicts of interest.

## Ethics Approval Statement

All samples were collected in accordance with relevant institutional Animal Ethics Committee (AEC) approvals and state permitting requirements. Collections in New South Wales were undertaken according to approvals from the University of New South Wales Animal Care and Ethics Committee (127B), Macquarie University AEC (2014/057-42), the NSW Office of Environment and Heritage (021028/02), Taronga Conservation Society Australia AEC (4a/04/21), and NSW NPWS Scientific Licenses (SL100821, SL100668, S11771). Victorian samples were collected according to Deakin University AEC (27/2004, B34-2022) and state permits (10010709, 10011301, 10002899, 10003981, 10006666, 10007858). Tasmanian samples were collected under University of Tasmania AEC approval (23764) and state permits (FA23016, FA21015, FA22262, ES23017, ES21179, FA05228, 7339/05). South Australian samples were collected under Flinders University AEC approvals (E388, E449, BIOL4825) and state permits (Y26040, 10/2003, Q24796, 137). Western Australian samples were collected according to Murdoch University AEC (RW3345/21) and state permits (FO25000631, SF004716, CE000772).

## Data Accessibility and Benefit-Sharing

Raw sequence data generated and analysed for this project are available on the NCBI Sequence Read Archive (SRA) database under accession number PRJNA1541615. Code will be made available at https://github.com/l-ockert/little-penguin-population-genetics upon publication. Data will be released upon publication, but a reviewer link is accessible prior: <u>PRJNA1541615</u>.

## Author Contributions

LAR, WBS, and JD conceptualised the study. LEO, JD, BJP, CPB, MRW, DCN, EKC, JPYA, MJL, and DRS supplied samples for this project. LEO conducted laboratory work and formal analyses. LEO drafted the manuscript with input from LAR, WBS, and JD. All authors provided feedback on the manuscript and approved the final version before submission.

## Notes

### Competing Interest Statement

The authors have declared no competing interest.

