## Supplementary Material for "High connectivity of an iconic seabird species may buffer genetic consequences of population declines"

**Supplementary Tables**

**Table S1.** Summary of samples collected for use in this study, per sampling site. Sampling details were combined when collected in consecutive years and stored/extracted using the same method. See Figure 1 in main text for sample locations, with numbers corresponding to the number before each site in the 'Location (State)' column. Further details of protocols used for DNA extraction and bioinformatic sexing are outlined in the methods section. For storage and extraction methods of archived DNA, see Burridge et al. (2015). Previous analysis refers to previous genetic analysis.

| Location (State) | **Year(s)** | **Males** | **Females** | **Tissue Type** | **Storage Method** | **DNA Extraction Method** | **Previous Analysis**[1] |
|---|---|---|---|---|---|---|---|
| *Known Provenance* | | | | | | | |
| 1. Broughton Island (NSW) | 2013 | 4 | 6 | Blood | Longmire's Buffer | DNeasy Blood and Tissue Kit (Qiagen) | Vardeh et al. (2023) |
| | 2024-25 | 15 | 11 | Blood | Whatman® Grade 5 qualitative filter papers | Puregene Tissue Kit (Qiagen) | None |
| 2. Cabbage Tree Island (NSW) | 2006 | 3 | 3 | Archived DNA | | | Peucker et al. (2009) |
| | 2012 | 13 | 18 | Blood | Longmire's Buffer | DNeasy Blood and Tissue Kit (Qiagen) | Vardeh et al. (2023) |
| | 2024 | 6 | 6 | Blood | QIAcard FTA Micro | Puregene Tissue Kit (Qiagen) | None |
| 3. Lion Island (NSW) | 2007 | 0 | 4 | Archived DNA | | | Burridge (2020); Peucker et al. (2009) |
| | 2012 | 13 | 17 | Blood | Longmire's Buffer | DNeasy Blood and Tissue Kit (Qiagen) | Vardeh et al. (2023) |

| | | | | | | | |
|---|---|---|---|---|---|---|---|
| | 2023-24 | 4 | 3 | Blood | Whatman® Grade 5 qualitative filter papers | Puregene Tissue Kit (Qiagen) | None |
| 4. Manly (NSW) | 2012 | 15 | 8 | Blood | Longmire's Buffer | DNeasy Blood and Tissue Kit (Qiagen) | Vardeh et al. (2023) |
| | 2023 | 0 | 1 | Blood | Whatman® Grade 5 qualitative filter papers | Puregene Tissue Kit (Qiagen) | None |
| | 2023 | 0 | 1 | Kidney | Frozen | Puregene Tissue Kit (Qiagen) | None |
| 5. Five Islands (NSW) | 2012 | 9 | 10 | Blood | Longmire's Buffer | DNeasy Blood and Tissue Kit (Qiagen) | Vardeh et al. (2023) |
| | 2024 | 5 | 5 | Blood | QIAcard FTA Micro | Puregene Tissue Kit (Qiagen) | None |
| 6. Bowen Island (NSW) | 2012 | 15 | 15 | Blood | Longmire's Buffer | DNeasy Blood and Tissue Kit (Qiagen) | Vardeh et al. (2023) |
| 7. Brush Island (NSW) | 2012 | 10 | 9 | Blood | Longmire's Buffer | DNeasy Blood and Tissue Kit (Qiagen) | Vardeh et al. (2023) |
| | 2024 | 8 | 12 | Blood | QIAcard FTA Micro | Puregene Tissue Kit (Qiagen) | None |
| 8. Snapper Island (NSW) | 2023-24 | 17 | 16 | Blood | Whatman® Grade 5 qualitative filter papers | Puregene Tissue Kit (Qiagen) | None |
| 9. Tollgate Islands (NSW) | 2023-24 | 18 | 16 | Blood | Whatman® Grade 5 qualitative filter papers | Puregene Tissue Kit (Qiagen) | None |

| | | | | | | | |
|---|---|---|---|---|---|---|---|
| 10. Montague Island (Barunguba) (NSW) | 2006-07 | 3 | 4 | Archived DNA | | | Peucker et al. (2009) |
| | 2012 | 17 | 12 | Blood | Longmire's Buffer | DNeasy Blood and Tissue Kit (Qiagen) | Vardeh et al. (2023) |
| | 2022-23 | 15 | 13 | Blood | Whatman® Grade 5 qualitative filter papers | Puregene Tissue Kit (Qiagen) | None |
| 11. Gabo Island (VIC) | 2025 | 10 | 10 | Blood | QIAcard FTA Micro | Puregene Tissue Kit (Qiagen) | None |
| 12. Furneaux Islands (TAS) | 2021-22 | 2 | 2 | Blood | Frozen | Puregene Tissue Kit (Qiagen) | None |
| 13. Bicheno (TAS) | 2021-22 | 4 | 2 | Blood | Frozen | Puregene Tissue Kit (Qiagen) | None |
| 14. Diamond Island (TAS) | 2021-22 | 3 | 6 | Blood | Frozen | Puregene Tissue Kit (Qiagen) | None |
| 15. Coswell Beach (TAS) | 2021-22 | 1 | 4 | Blood | Frozen | Puregene Tissue Kit (Qiagen) | None |
| 16. Little Christmas Island (TAS) | 2021-22 | 1 | 10 | Blood | Frozen | Puregene Tissue Kit (Qiagen) | None |
| 17. Pirates Bay (TAS) | 2021-22 | 2 | 3 | Blood | Frozen | Puregene Tissue Kit (Qiagen) | None |
| 18. Wedge Island (TAS) | 2021 | 1 | 3 | Blood | Frozen | Puregene Tissue Kit (Qiagen) | None |

| | | | | | | | |
|---|---|---|---|---|---|---|---|
| 19. Bruny Island (TAS) | 2006 | 5 | 4 | Archived DNA | | | Burridge et al. (2015); Peucker et al. (2009) |
| 20. Boronia Beach (TAS) | 2021-22 | 6 | 0 | Blood | Frozen | Puregene Tissue Kit (Qiagen) | None |
| 21. Red Chapel Beach (TAS) | 2021-22 | 2 | 2 | Blood | Frozen | Puregene Tissue Kit (Qiagen) | None |
| 22. Macquarie Heads (TAS) | 2022 | 4 | 3 | Blood | Frozen | Puregene Tissue Kit (Qiagen) | None |
| 23. Stanley (TAS) | 2021 | 2 | 3 | Blood | Frozen | Puregene Tissue Kit (Qiagen) | None |
| 24. Burnie (TAS) | 2021-23 | 12 | 11 | Blood | Frozen | Puregene Tissue Kit (Qiagen) | None |
| 25. King Island (TAS) | 2022 | 9 | 9 | Blood | Frozen | Puregene Tissue Kit (Qiagen) | None |
| 26. Seal Island (VIC) | 2025 | 11 | 9 | Blood | QIAcard FTA Micro | Puregene Tissue Kit (Qiagen) | None |
| 27. Kanowna Island (VIC) | 2025 | 9 | 11 | Blood | QIAcard FTA Micro | Puregene Tissue Kit (Qiagen) | None |
| 28. Phillip Island (VIC) | 2003-04 | 8 | 6 | Archived DNA | | | Burridge et al. (2015); Overeem et al. (2008); Peucker et al. (2009) |
| | 2017-24 | 13 | 12 | Muscle | Ethanol | Puregene Tissue Kit (Qiagen) | None |

| | | | | | | | |
|---|---|---|---|---|---|---|---|
| 29. London Bridge (VIC) | 2003-04 | 8 | 6 | Archived DNA | | | Burridge et al. (2015); Overeem et al. (2008); Peucker et al. (2009) |
| 30. Middle Island (VIC) | 2003 | 5 | 8 | Archived DNA | | | Burridge et al. (2015); Overeem et al. (2008); Peucker et al. (2009) |
| 31. Granite Island (SA) | 2003 | 7 | 4 | Archived DNA | | | Burridge et al. (2015); Overeem et al. (2008); Peucker et al. (2009) |
| | 2013 | 1 | 4 | Blood | QIAcard FTA Classic | Puregene Tissue Kit (Qiagen) | Colombelli-Négrel et al. (2020) |
| 32. West Island (SA) | 2004 | 11 | 7 | Archived DNA | | | Burridge et al. (2015); Peucker et al. (2009) |
| 33. Antechamber Bay (SA) | 2013-14 | 10 | 8 | Blood | QIAcard FTA Classic | Puregene Tissue Kit (Qiagen) | Colombelli-Négrel et al. (2020) |
| 34. Kingscote (SA) | 2013-15 | 7 | 3 | Blood | QIAcard FTA Classic | Puregene Tissue Kit (Qiagen) | Colombelli-Négrel et al. (2020) |
| 35. Emu Bay (SA) | 2013-14 | 10 | 7 | Blood | QIAcard FTA Classic | Puregene Tissue Kit (Qiagen) | Colombelli-Négrel et al. (2020) |
| 36. Troubridge Island (SA) | 2013-15 | 19 | 14 | Blood | QIAcard FTA Classic | Puregene Tissue Kit (Qiagen) | Colombelli-Négrel et al. (2020) |
| 37. Spencer Gulf (SA) | 2006 | 13 | 8 | Archived DNA | | | Burridge et al. (2015); Peucker et al. (2009) |

| | | | | | | | |
|---|---|---|---|---|---|---|---|
| 38. Pearson Island (SA) | 2006 | 8 | 10 | Archived DNA | | | Burridge et al. (2015); Peucker et al. (2009) |
| 39. St Peter Island (SA) | 2006 | 4 | 2 | Archived DNA | | | Peucker et al. (2009) |
| 40. Cheyne Island (WA) | 2005 | 14 | 9 | Archived DNA | | | Burridge et al. (2015); Peucker et al. (2009) |
| 41. Breaksea Island (WA) | 2007 | 3 | 4 | Archived DNA | | | Peucker et al. (2009) |
| 42. Penguin Island (WA) | 2005 | 2 | 1 | Archived DNA | | | Burridge et al. (2015); Peucker et al. (2009) |
| | 2024 | 6 | 4 | Blood | Whatman® Grade 5 qualitative filter papers | Puregene Tissue Kit (Qiagen) | None |
| *Unknown Origin and Vagrants*[2] | | | | | | | |
| Ichaboe Island, Namibia | 2005 | 0 | 1 | Archived DNA | | | Peucker et al. (2009) |
| Palm Beach (NSW) | 2022 | 1 | 0 | Blood | QIAcard FTA Classic | Puregene Tissue Kit (Qiagen) | None |
| Barlings Beach (NSW) | 2023 | 1 | 2 | Kidney | Frozen | Puregene Tissue Kit (Qiagen) | None |
| Coogee Beach (NSW) | 2024 | 0 | 1 | Blood | Whatman® Grade 5 qualitative filter papers | Puregene Tissue Kit (Qiagen) | None |
| Goolwa Barrage (SA) | 2024 | 1 | 0 | Muscle | Frozen | Puregene Tissue Kit (Qiagen) | None |

| | | | | | | | |
|---|---|---|---|---|---|---|---|
| Hindmarsh River (SA) | 2024 | 0 | 2 | Muscle | Frozen | Puregene Tissue Kit (Qiagen) | None |
| Encounter Bay Croquet Club (SA) | 2024 | 0 | 1 | Muscle | Frozen | Puregene Tissue Kit (Qiagen) | None |
| Hayborough (SA) | 2024 | 0 | 1 | Muscle | Frozen | Puregene Tissue Kit (Qiagen) | None |
| Middleton (SA) | 2024 | 4 | 0 | Muscle | Frozen | Puregene Tissue Kit (Qiagen) | None |
| Albany (WA) | 2025 | 1 | 2 | Skin | Ethanol | Puregene Tissue Kit (Qiagen) | None |
| Esperance (WA) | 2025 | 0 | 4 | Skin | Ethanol | Puregene Tissue Kit (Qiagen) | None |

[1] Previous studies analysed partially overlapping samples from these sampling efforts; not all samples were available or suitable for both studies. Note that varying genetic markers were used in each study, including microsatellites (Peucker et al., 2009), microsatellites and mitochondrial DNA (Burridge et al., 2015; Overeem et al., 2008; Vardeh et al., 2023), and single nucleotide polymorphisms (Colombelli-Négrel et al., 2020).

[2] Location refers to site where beach-washed/injured individuals were recovered from and is not necessarily reflective of their colony of origin.

**Table S2.** Comparison of the proportion of trimmed, filtered, and replicate-merged DArTseq reads (ranging from 60-100 bp in length) that mapped to four reference genomes using the Burrows Wheeler Alignmener algorithms, *BWA-MEM* and *BWA-ALN*. A subset of 50 samples representing the temporal and spatial range of the dataset was used to reduce computational demands associated with the full dataset. Mapping rates were determined using SAMtools *flagstat*.

| **Reference Genome** | | | | **Mean Mapping Rate (%)** | |
|---|---|---|---|---|---|
| **Assembly** | **Species** | **Level** | **Sex** | ***BWA-MEM*** | ***BWA-ALN*** |
| BGI_Enov.V1 (Pan et al., 2019) | *Eudyptula minor novaehollandiae* | Scaffold | Not reported | 96.83 | 51.08 |
| DNAZoo (DNA Zoo, 2024) | *Eudyptula minor* | Scaffold | Not reported | 96.53 | 50.59 |
| bSphHub1.pri (Rhie et al., 2021) | *Spheniscus humboldti* | Chromosome | Female | 95.18 | 46.68 |
| bSphDem1.hap1 (Rhie et al., 2021) | *Spheniscus demersus* | Chromosome | Male | 95.02 | 46.65 |

**Table S3.** Comparison of two common variant calling pipelines; BCFtools *mpileup/call* and Stacks *gstacks/populations*. Variant calling was conducted using reads aligned to the *Spheniscus humboldti* reference genome with *BWA-MEM*. Variant counts were reported using BCFtools. Per-site and per-individual missingness was calculated using VCFtools *missing-site* and *missing-indv* flags, respectively. Both datasets were filtered with VCFtools under the same low-stringency conditions, retaining biallelic SNPs with a minor allele count ≥ 2 that were present in at least 50% of individuals, had a minimum depth of 5X, and a mean depth ≤ 20X.

| | **Unfiltered Dataset** | | | **Filtered Dataset** | | |
|---|---|---|---|---|---|---|
| **Variant Caller** | **Number of Variants** | **Mean Missingness per Indv (%)** | **Mean Depth per Indv (X)** | **Number of Variants** | **Mean Missingness per Indv (%)** | **Mean Depth per Indv (X)** |
| *BCFtools* | 1,060,463 | 65.26 | 2.78 | 14,655 | 25.10 | 10.91 |
| *Stacks* | 195,622 | 42.35 | 7.38 | 15,299 | 26.40 | 11.69 |

**Table S4.** Number of SNPs and individuals retained after initial filtering steps, and the thresholds applied to generate datasets for different analyses. Monomorphic loci were filtered when individuals were removed.

| Initial filtering step | SNPs retained | Individuals retained |
|---|---|---|
| Initial stacks calling | 122,573 | 1,012 |
| Read depth, 3-26X | 12,341 | 1,012 |
| Merging sequencing replicates | 12,341 | 842 |
| Difference in coverage ≤ 70% | 11,405 | 842 |
| Reproducibility ≥ 90% | 11,316 | 842 |
| Heterozygosity ≤ 70% | 11,248 | 842 |
| High genotype similarity | 10,811 | 838 |
| Sex-linked loci | 10,243 | 838 |
| Individual call rate ≤ 50% | 10,103 | 829 |
| Relatedness ≥ 0.4 | 9,967 | 796 |
| Excluding unknown provenance[1] | 9,898 | 773 |

**Filtering thresholds applied for analyses**

| | Dataset 1. Structure | Dataset 2. Gene Flow | Dataset 3. Diversity | Dataset 4. Provenance |
|---|---|---|---|---|
| Minor allele count | ≤ 3 | N/A | ≤ 2 | ≤ 3 |
| Minor allele frequency | N/A | ≥ 0.05 | N/A | N/A |
| Loci call rate | ≥ 60% | ≥ 98% | ≥ 80% | ≥ 80% |
| Individual call rate | ≥ 60% | ≥ 80% | ≥ 80% | ≥ 80% |
| Total SNPs | 5,982 | 1,459 | 5,615 | 5,322 |
| Total individuals | 772 | 772 | 766 | 788 |

[1] This filtering step was excluded when generating the dataset for provenance analyses.

**Table S5.** Genetic differentiation between the four genetic groups of little penguins as previously identified by admixture analyses. $F_{ST}$ is represented below the diagonal while Mutual Information is represented above the diagonal. Estimates include ±95% confidence intervals. NNS = Northern New South Wales, EAC = East Australian Coast, BST = Bass Strait and Tasmania, and SWA = Southern and Western Australia.

| | **NNS** | **EAC** | **BST** | **SWA** |
|---|---|---|---|---|
| **NNS** | | 0.0033 (±0.0354) | 0.0041 (±0.0332) | 0.0100 (±0.0393) |
| **EAC** | 0.0055 (±0.0005) | | 0.0012 (±0.0231) | 0.0072 (±0.0269) |
| **BST** | 0.0077 (±0.0006) | 0.0032 (±0.0003) | | 0.0080 (±0.0317) |
| **SWA** | 0.0252 (±0.0016) | 0.0217 (±0.0011) | 0.0201 (±0.0011) | |

**Table S6.** Rarefied private allele counts within each group across three time periods. Values were rarefied to the time point with the smallest sample size within each group (denoted in parenthesis in the 'group' column), and confidence intervals (denoted in parenthesis next to the mean) were calculated with 1000 bootstraps. Calculations for SWA excluded samples from Western Australia due to uneven temporal sampling, and therefore only include samples from South Australia.

| | **Time Period** | | |
|---|---|---|---|
| **Group** | A | B | C |
| NNS ($n$ = 6) | 282.00 (252.00 – 315.02) | 285.25 (242.00 – 331.00) | 298.26 (260.00 – 337.00) |
| EAC ($n$ = 9) | 246.54 (223.00 – 272.02) | 295.49 (261.00 – 329.00) | 279.79 (246.00 – 313.00) |
| BST ($n$ = 48) | 395.48 (368.00 – 425.00) | N/A | 369.09 (347.00 – 391.02) |
| SWA ($n$ = 67) | 446.54 (442.00 – 456.00) | 346.84 (337.00 – 350.00) | N/A |

### **Supplementary Figures**

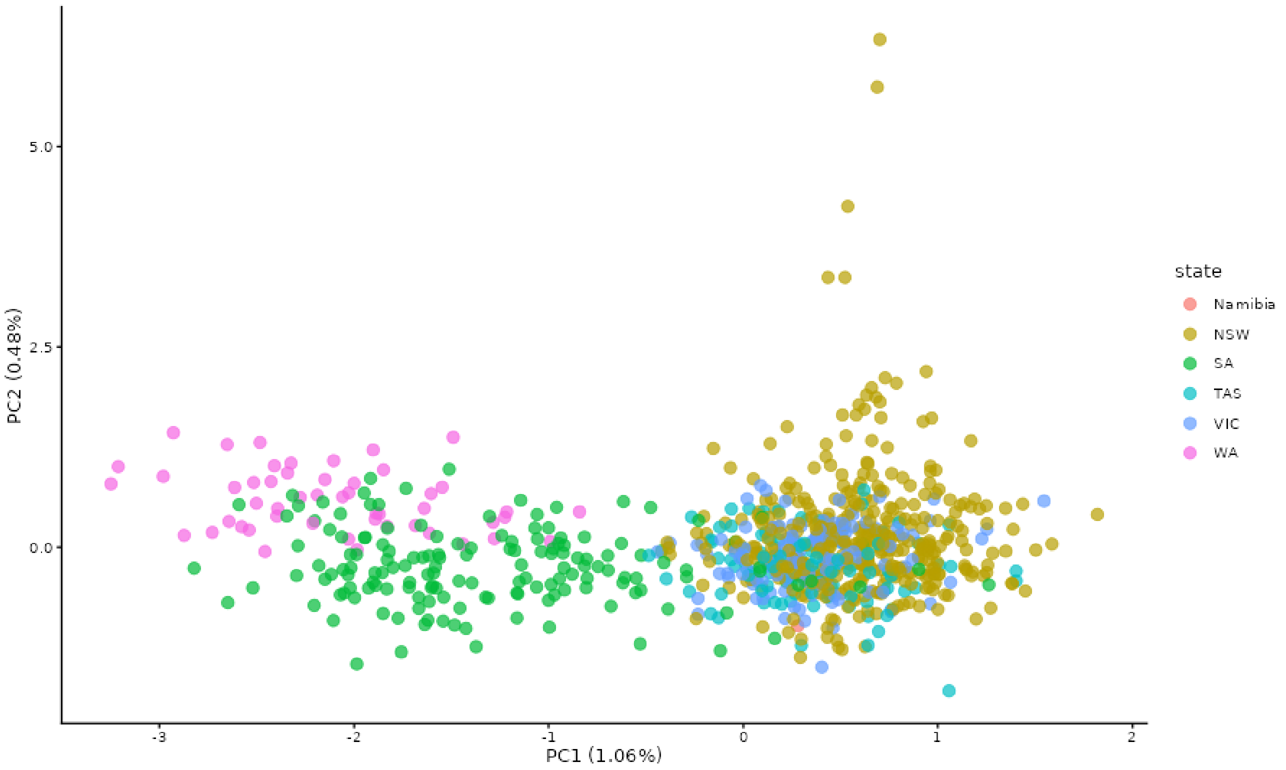


**Figure S1.** Initial PCA plot used to determine broad groupings for Stacks *populations* module. Dataset was generated using Stacks to call variants (without populations specified in the population map file), and filtering using VCFtools based on initial thresholds recommended by Hemstrom et al. (2024): *--minDP* 5, *--maxmeanDP* 10, *--max-missing* 0.6, *--mac* 3, *--hwe* 1e06, and removing individuals with >50% missing data. PCA was generated using PLINK.

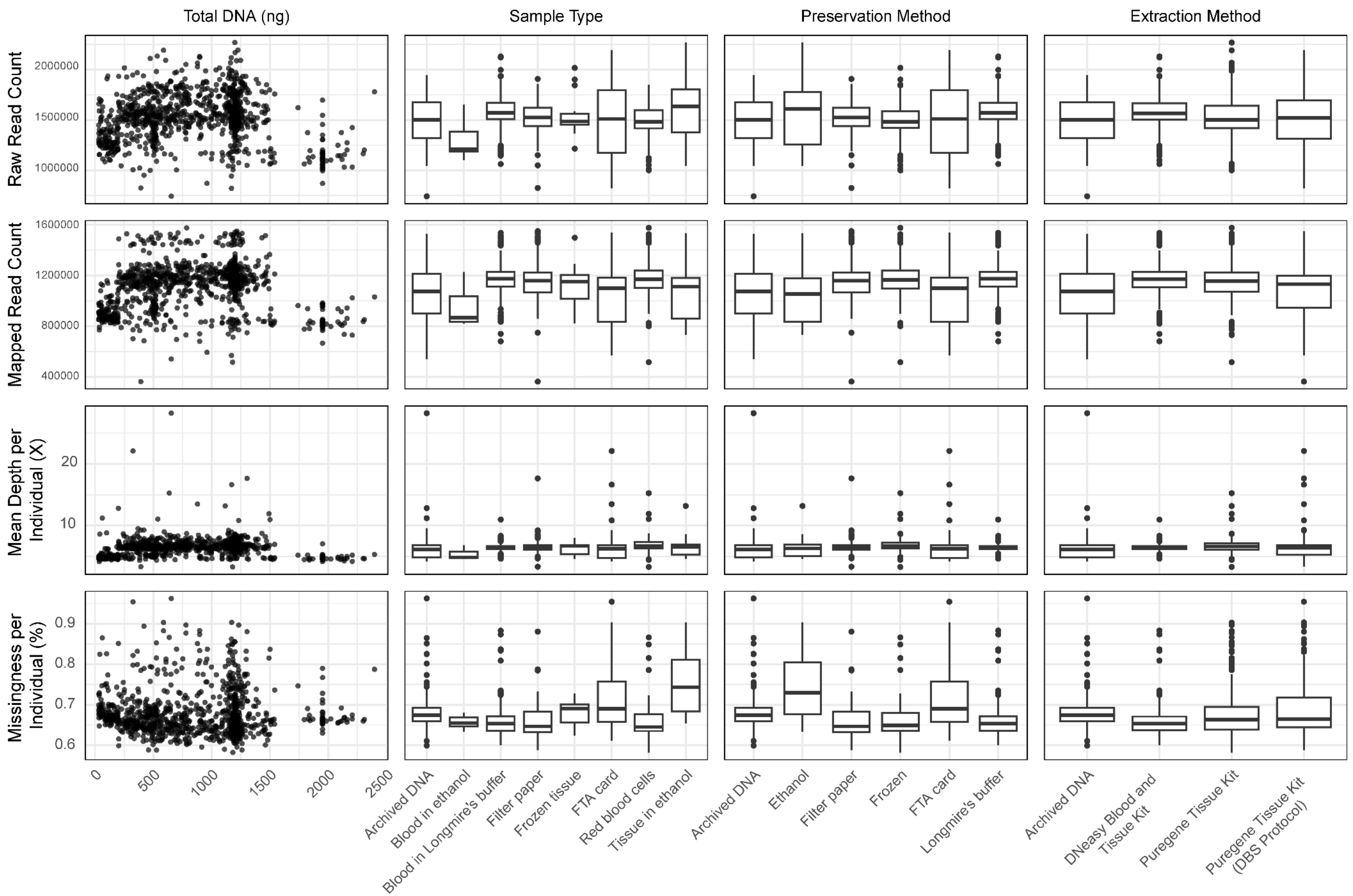


48 **Figure S2.** Comparison of sequencing performance across different sample characteristics. DNA quantity (Total DNA), sample type, sample preservation

method, and extraction method are compared against sequencing metrics raw read counts, mapped read counts, mean sequencing depth per individual (pre-filtering), and the proportion of missing sites per individual (pre-filtering).

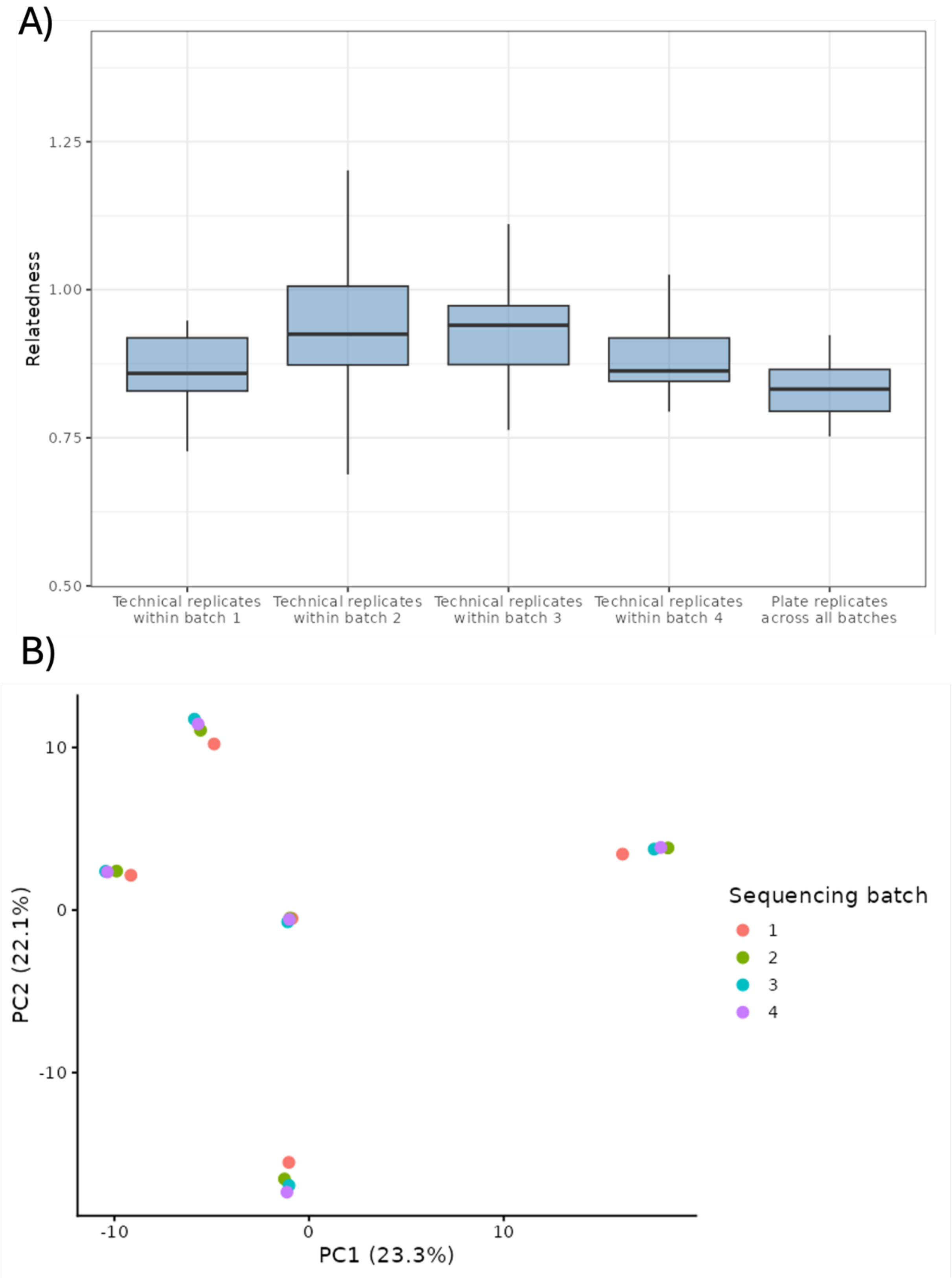


51 **Figure S3.** Consistency between four sequencing batches. (A) Pairwise $A_{JK}$ relatedness
52 between technical replicates and their corresponding original sequence file, and between five
53 plate replicates that were included in all four batches. (B) PCoA of the five plate replicates,
54 where each colour corresponds to a different sequencing batch.

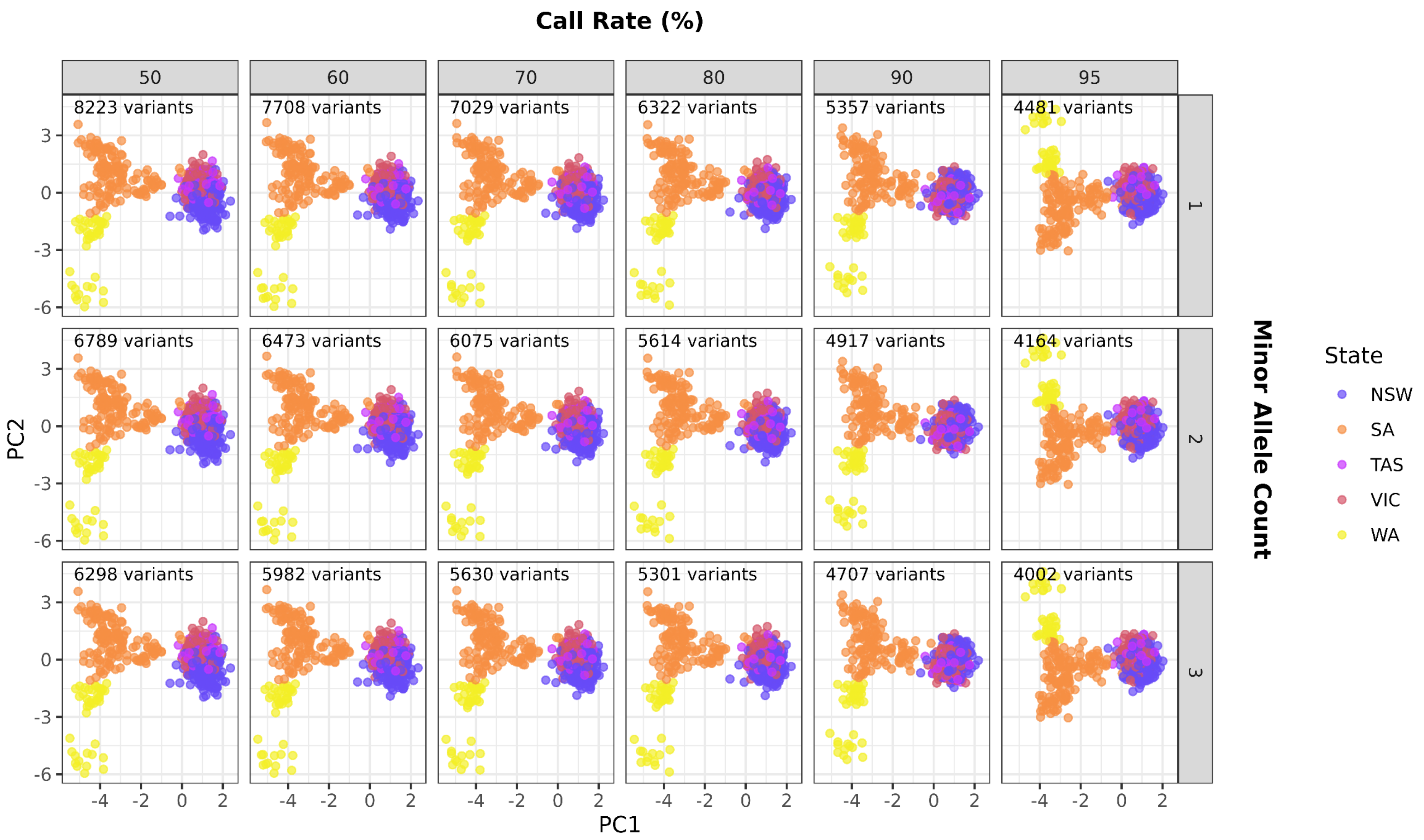
Call Rate (%)
50
60
70
80
90
95
8223 variants
7708 variants
7029 variants
6322 variants
5357 variants
4481 variants
6789 variants
6473 variants
6075 variants
5614 variants
4917 variants
4164 variants
6298 variants
5982 variants
5630 variants
5301 variants
4707 variants
4002 variants
1
2
3
Minor Allele Count
PC2
PC1
State
NSW
SA
TAS
VIC
WA

55

**Figure S4.** Comparison of call rate (CR) and minor allele count (MAC) filtering thresholds. Principal Coordinates Analyses (PCoA) were plotted to visualise the impact of filtering on population structure, with the number of variant loci retained in each dataset listed in the top right corner. Details of initial variant calling can be found in the main methods section.

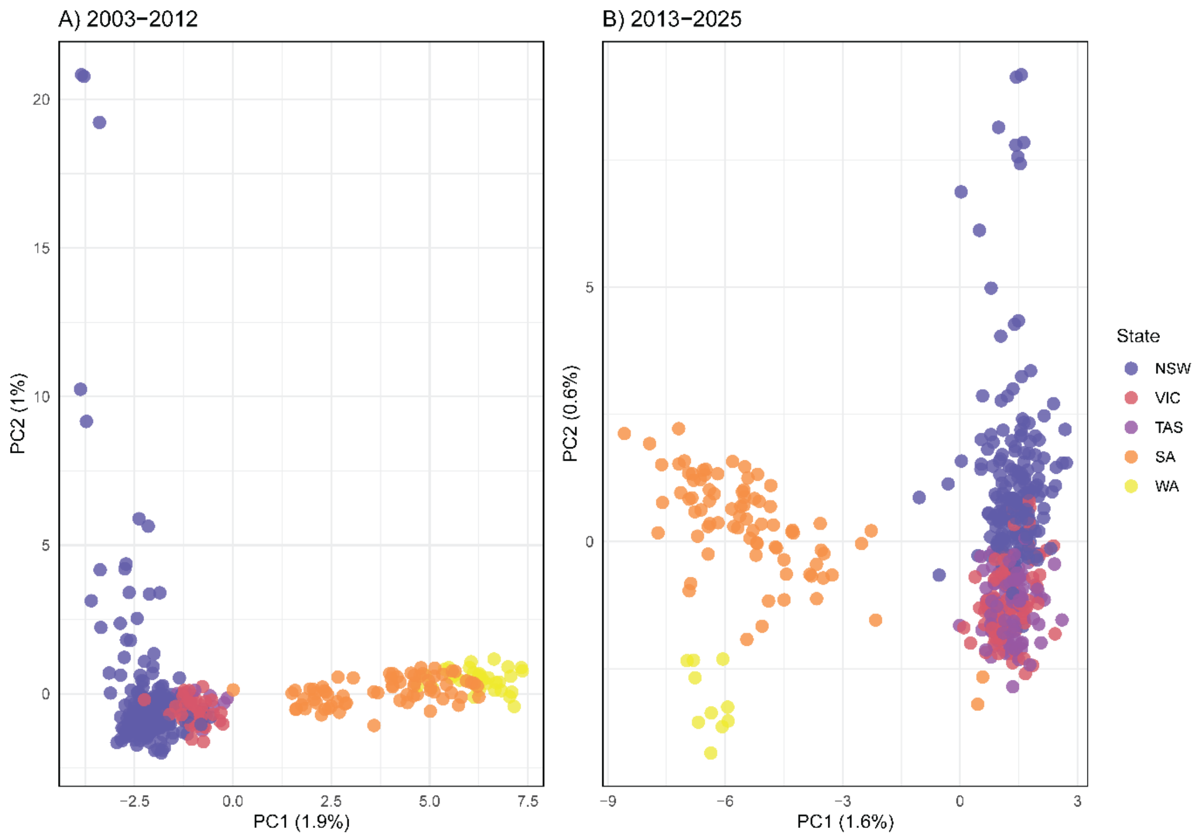


**Figure S5.** Principal Coordinates Analysis (PCoA) plot of all samples of known origin, collected between A) 2003-2012 and B) 2013-2025. Points are coloured by state, where NSW = New South Wales, VIC = Victoria, TAS = Tasmania, SA = South Australia, and WA = Western Australia. Note that the same colonies were not necessarily sampled during both time periods.

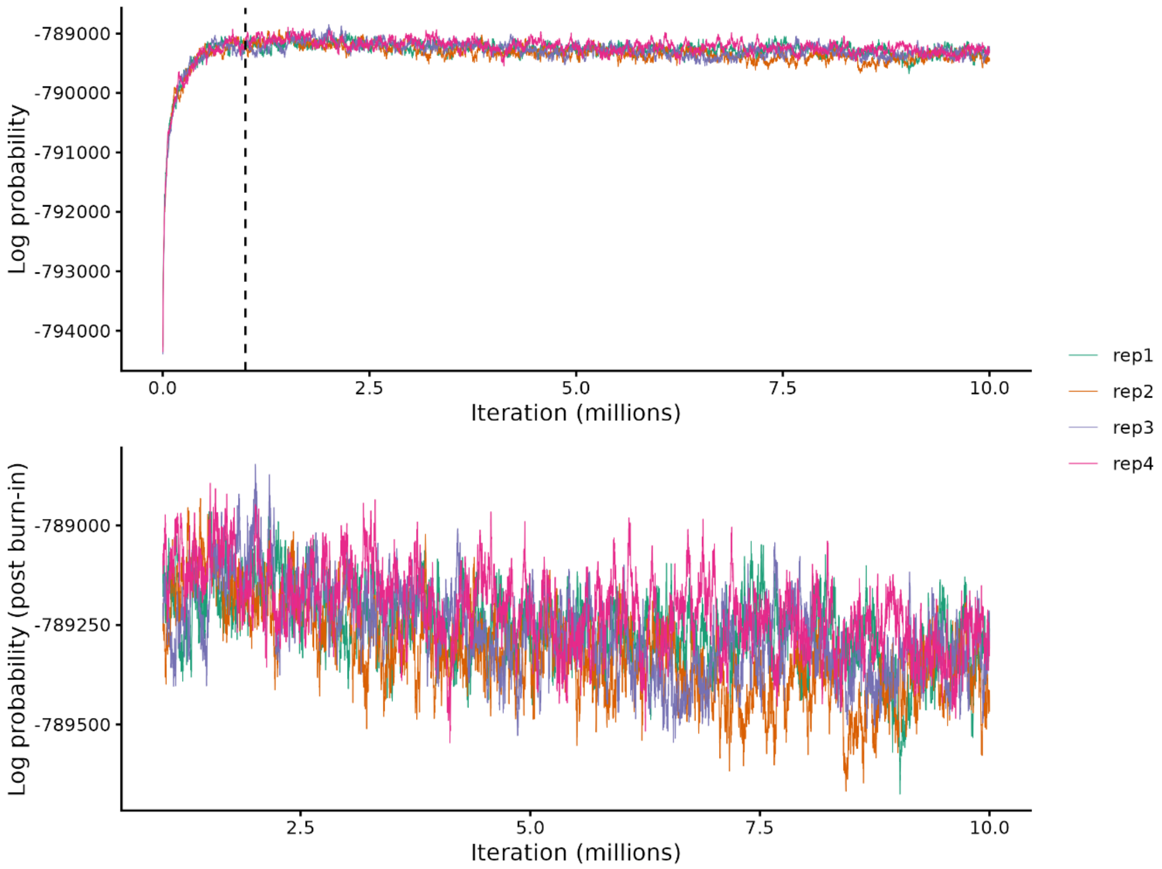


**Figure S6.** MCMC trace of the log-probability for four independent BayesAss runs. Each run used a different random seed and involved 5,000,000 iterations, a burn-in of 1,000,000 iterations, and a sampling interval of 1,000. The dashed line in the top plot indicates the 1,000,000 burn-in iterations and the bottom plot indicates the post burn-in sampling.

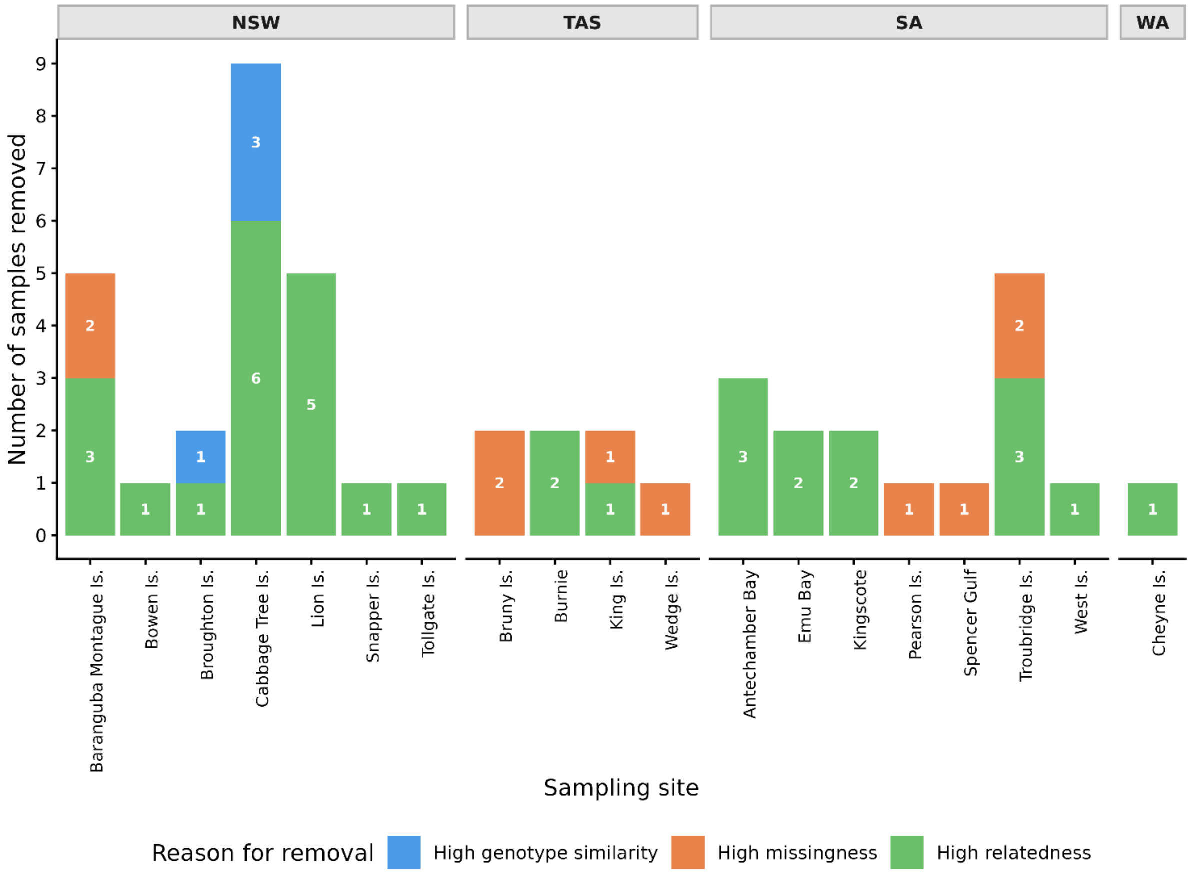


**Figure S7.** The number of individuals removed, per sampling site, due to either being identified as a duplicate sample (genotype similarity > 99%), exhibiting a high proportion of missing data (> 60%), or being highly related to other individuals (*R* > 0.4). Duplicated and highly related individuals are identified pairwise, hence, one individual was removed from each pair while the other was retained.

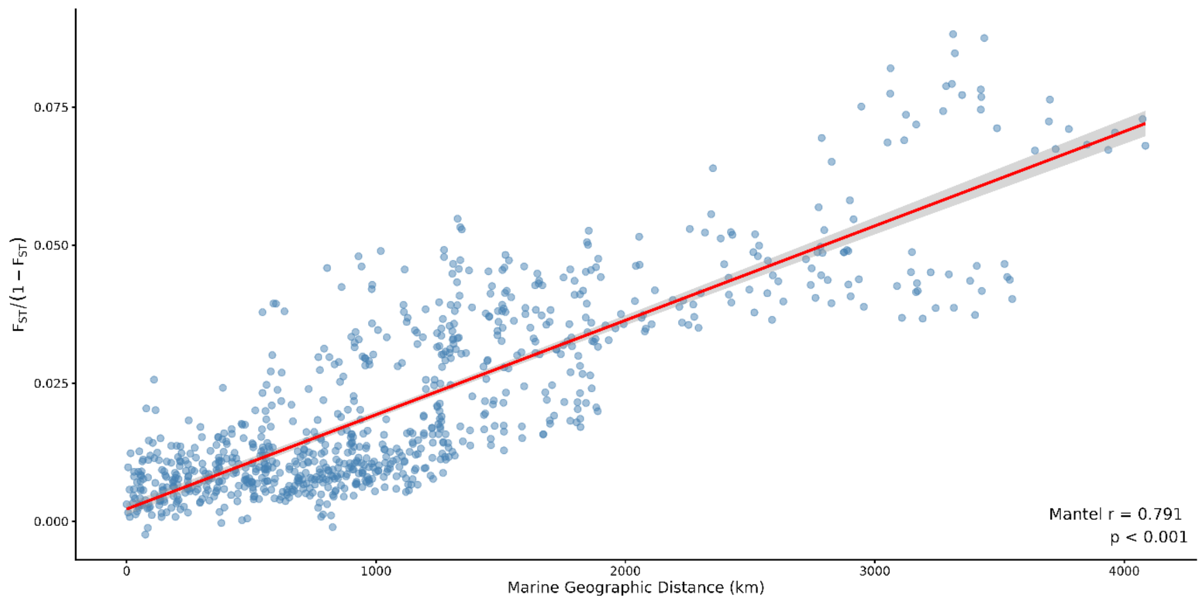


**Figure S8.** Correlation between linearised genetic distance ($F_{ST}$) and marine geographic distance (km), demonstrating little penguins display a pattern of isolation by distance. The red line indicates a linear relationship with a confidence band shown in grey.

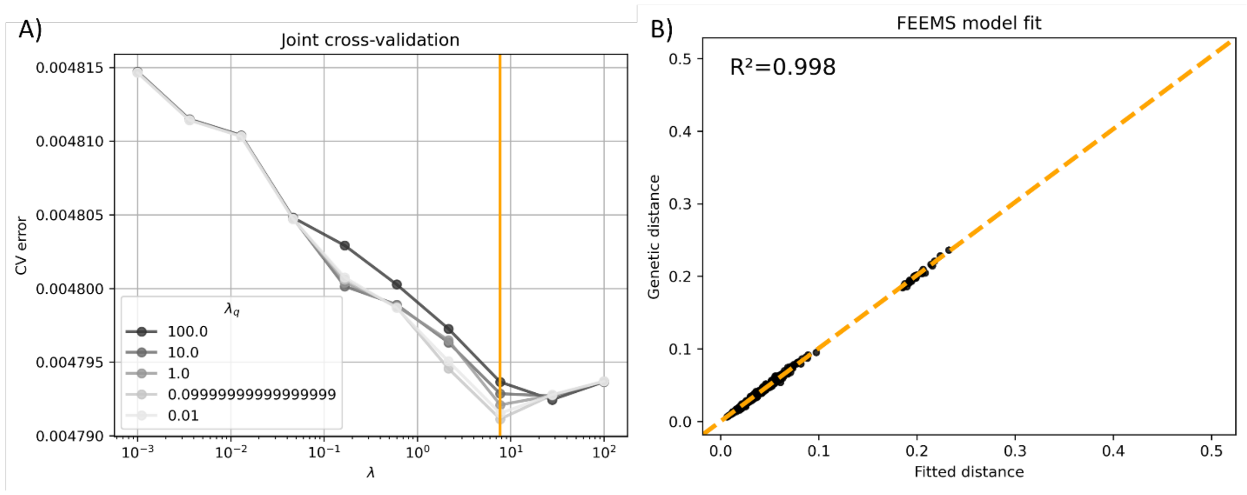


**Figure S9.** FEEMS model selection and fit. (A) Joint cross-validation error across a range of smoothing parameters (λ and λ_q). (B) FEEMS model fit showing the relationship between fitted and observed pairwise genetic distances across all sample pairs.

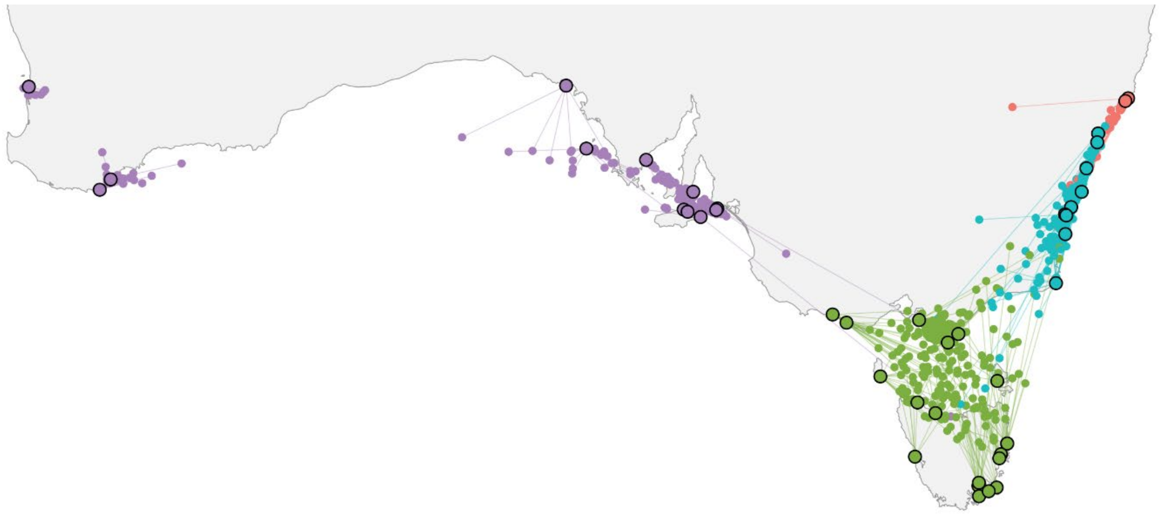

**Figure S10.** Accuracy of locator geographic predictions assessed by 10-fold cross-validation, based on predicted location of all 766 known provenance samples. Black outlined circles indicate the coordinates of the sample location, while non-outlined points indicate the predicted coordinates, and lines indicate the true location of the predicted coordinates. Colours indicate the genetic grouping that contains each sampled colony and predicted assignment, where red = NNS, blue = EAC, green = BST, and purple = SWA.

Rhie, A., McCarthy, S. A., Fedrigo, O., Damas, J., Formenti, G., Koren, S., Uliano-Silva, M., Chow, W., Fungtammasan, A., Kim, J., Lee, C., Ko, B. J., Chaisson, M., Gedman, G.

L., Cantin, L. J., Thibaud-Nissen, F., Haggerty, L., Bista, I., Smith, M., Haase, B., Mountcastle, J., Winkler, S., Paez, S., Howard, J., Vernes, S. C., Lama, T. M., Grutzner, F., Warren, W. C., Balakrishnan, C. N., Burt, D., George, J. M., Biegler, M. T., Iorns, D., Digby, A., Eason, D., Robertson, B., Edwards, T., Wilkinson, M., Turner, G., Meyer, A., Kautt, A. F., Franchini, P., Detrich, H. W., 3rd, Svardal, H., Wagner, M., Naylor, G. J. P., Pippel, M., Malinsky, M., Mooney, M., Simbirsky, M., Hannigan, B. T., Pesout, T., Houck, M., Misuraca, A., Kingan, S. B., Hall, R., Kronenberg, Z., Sovic, I., Dunn, C., Ning, Z., Hastie, A., Lee, J., Selvaraj, S., Green, R. E., Putnam, N. H., Gut, I., Ghurye, J., Garrison, E., Sims, Y., Collins, J., Pelan, S., Torrance, J., Tracey, A., Wood, J., Dagnew, R. E., Guan, D., London, S. E., Clayton, D. F., Mello, C. V., Friedrich, S. R., Lovell, P. V., Osipova, E., Al-Ajli, F. O., Secomandi, S., Kim, H., Theofanopoulou, C., Hiller, M., Zhou, Y., Harris, R. S., Makova, K. D., Medvedev, P., Hoffman, J., Masterson, P., Clark, K., Martin, F., Howe, K., Flicek, P., Walenz, B. P., Kwak, W., Clawson, H., Diekhans, M., Nassar, L., Paten, B., Kraus, R. H. S., Crawford, A. J., Gilbert, M. T. P., Zhang, G., Venkatesh, B., Murphy, R. W., Koepfli, K. P., Shapiro, B., Johnson, W. E., Di Palma, F., Marques-Bonet, T., Teeling, E. C., Warnow, T., Graves, J. M., Ryder, O. A., Haussler, D., O'Brien, S. J., Korlach, J., Lewin, H. A., Howe, K., Myers, E. W., Durbin, R., Phillippy, A. M., & Jarvis, E. D. (2021). Towards complete and error-free genome assemblies of all vertebrate species. *Nature, 592*(7856), 737–746. doi: 10.1038/s41586-021-03451-0

Vardeh, S., Sinclair, J., Cairns, K., Cannell, B., Rollins, L., & Sherwin, W. (2023). Genetic connectivity of little penguin colonies in Australia. *Authorea*. doi: 10.22541/au.168569955.57431501/v1
